# A representative of a ubiquitous bacterial lineage parasitically feeds on host RNA

**DOI:** 10.64898/2026.06.21.733656

**Authors:** Taiki Katayama, Naoki Hosogi, Xian-Ying Meng, Yoichi Kamagata, Hideyuki Tamaki, Masaru K. Nobu

**Author notes:** Corresponding authors: Taiki Katayama, Masaru K. Nobu. These authors contributed equally to this work (Taiki Katayama, Masaru K. Nobu).

## Abstract

Cellular metabolism is widely understood as an integrated network of redox reactions, energy conservation, and biosynthetic pathways. Here we show that across diverse prokaryotic lineages, loss of redox-associated functions is coupled with loss of nucleotide biosynthesis, raising a fundamental question of how such metabolically reduced organisms sustain cell growth. One of the most widespread and diverse lineages of prokaryotes—*Minisyncoccota* or Patescibacteriota—constitutes the majority of lineages exhibiting this pattern. To investigate how such organisms persist, we cultivated and characterized a representative of this lineage from a deep aquifer. The organism attaches to and penetrates growing bacterial host cells, and directly uptakes host RNA, concomitant with its depletion in the host. The metabolically reduced parasite cleaves host-derived RNA to directly supply cellular energy currencies and precursors for RNA/DNA synthesis, NTPs, without invoking canonical metabolic pathways. Codon usage in the parasite is complementary to that of its host, potentially minimizing translation of host-derived mRNA. Comparative genomics and phylogenetics indicate that these features are widespread and likely ancestral across the lineage. Exploitation of host RNA as a metabolic resource reveals a previously unrecognized metabolic strategy that demonstrates cellular metabolism can be sustained through direct utilization of informational macromolecules and thereby taps into an omnipresent energy reservoir, potentially supporting the environmental ubiquity of the metabolically reduced lineage.

## Introduction

Cellular metabolism is fundamentally organized around redox chemistry. The transfer of electrons from donors to acceptors drives energy conservation and provides the reducing power required for biosynthesis. This reliance on electron flow, energy conservation, and biosynthesis is widely regarded as a central organizing principle of cellular life.

Yet a substantial fraction of prokaryotic diversity appears to operate at the margins of this principle. Members of the *Minisyncoccota*^1^ (ICNP; corresponding to Patescibacteriota^2,3^ under the SeqCode and historically Candidate Phyla Radiation or CPR^4^)—a vast and ubiquitous bacterial radiation comprising a substantial fraction (8 to 15%) of bacterial diversity^4,5^—possess highly reduced genomes that lack many canonical metabolic pathways. Previous comparative analyses have also suggested that these genomes are depleted in proteins associated with redox metabolism^6^, raising a fundamental question: how do these organisms sustain the energetic and biosynthetic demands of cellular growth?

Here we report the cultivation and characterization of a bacterium that resolves this paradox through a metabolic strategy that departs from canonical redox-based metabolism. This organism grows as an ectoparasite on a bacterial host and directly exploits host-derived RNA, importing and degrading it to nucleotides that serve as central metabolic intermediates and, potentially, as energy carriers. Genomic, physiological, and imaging data collectively indicate that this RNA-centered metabolism operates with minimal reliance on canonical electron-transfer pathways.

Beyond revealing a previously unrecognized mode of host exploitation, these findings suggest that cellular metabolism can be sustained through direct assimilation of informational macromolecules, decoupling metabolism from canonical redox-coupled energy conservation. This expands the known repertoire of biological energy strategies and provides a mechanistic framework for understanding the ecological success of one of the most widespread bacterial radiations on Earth.

### Coupled depletion of redox metabolism and nucleotide biosynthesis

We examined the metabolic capacities of reduced microbial genomes across prokaryotic lineages and found a striking and previously unappreciated coupling between redox metabolism, including energy conservation via respiration, and *de novo* nucleotide biosynthesis: genomes depleted in one were consistently depleted in the other (Fig. 1a and 1b). This leaves a subset of organisms with minimal capacity for redox metabolism and nucleotide production, a pattern observed across diverse prokaryotes, raising the question of how such organisms sustain growth.

**Fig. 1.**
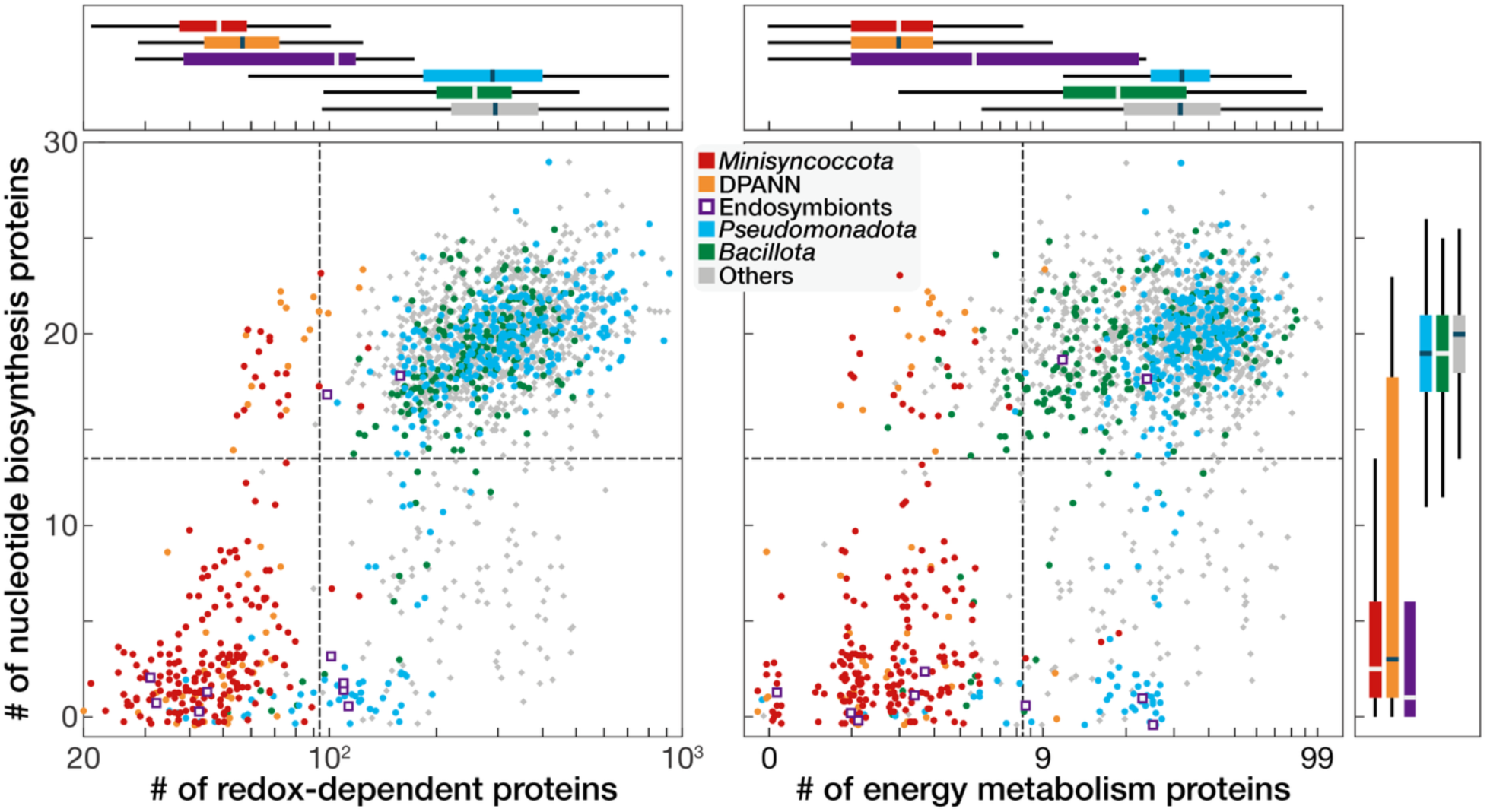
Distribution of functional protein categories across prokaryotic genomes. Each point represents the median protein count for genomes aggregated at the order level. Marginal boxplots show the distributions of the corresponding protein counts for each group. Dashed lines indicate the 1.5×IQR lower whisker values calculated from lineages excluding *Minisyncoccota*, DPANN, and endosymbiotic lineages. Endosymbiotic lineages include *Ca.* Babelota, *Chlamydiota*, *Borreliales*, *Rickettsiales*, *Paracaedibacterales*, *Caedimonadales*, and *Holosporales*. Small random jitter was added to point positions to reduce overplotting.

This pattern is observed across a limited set of host-associated lineages that employ distinct strategies to compensate for reduced metabolic capacity. Some intracellular parasites, including *Chlamydiota*, *Rickettsiales*, and *Caedimonadales*, acquire nucleotides directly from their hosts via ATP/ADP translocases^7–12^ while retaining components of redox metabolism. Others, such as *Mycoplasmatales* and *Borreliales*, are further depleted in redox proteins and respiratory pathways but sustain growth through substrate-level metabolism of host-derived sugars or amino acids^13,14^. Obligate intracellular lineages, including members of *Ca.* Babelota and *Holosporales*, exhibit further dependence on host cytoplasmic resources. In contrast, members of *Minisyncoccota*/Patescibacteriota, which constitute the majority of lineages exhibiting reduced metabolic capacities, are generally observed to attach to, rather than invade, their hosts and lack identifiable energy metabolisms and nucleotide transport systems. These features indicate a previously unrecognized mode of host exploitation that may underlie a substantial fraction of metabolically reduced microbial diversity.

### A cultivated representative acquires host RNA

To investigate the biological basis of this reduced metabolic state, we sought to experimentally characterize a representative organism from this lineage. After 9 years of cultivation (see Supplementary Note 1), we obtained a pure co-culture consisting of ultrasmall cells (strain OT8) and filamentous cells (strain Mc4) from a deep anoxic aquifer in natural-gas deposits^15^. Phylogenetic analysis placed strain OT8 in the *Minisyncoccota*/Patescibacteriota class *Ca*. Microgenomatia (formerly OP11), a lineage that has eluded cultivation since its first detection 28 years ago^16^ (Supplementary Fig. S1). Its host, strain Mc4, was affiliated with the family *Aggregatilineaceae* within the phylum *Chloroflexota* (Supplementary Fig. S1).

Strain OT8 was consistently observed attached to Mc4 cells (Fig. 2a, b), suppressed host growth, proliferating only in the presence of actively growing Mc4 cells (Fig. 2c, d), and preferentially localized to host cells’ elongation sites (Supplementary Fig. S2). OT8 failed to grow in the presence non-growing Mc4 cells or in the absence of Mc4, or in the presence of growing cells of a close Mc4 relative species (Supplementary Fig. S3, Fig. S4 and Note 2), indicating that OT8 is an obligate, host-specific ectoparasite. Transmission-electron and cryo-electron microscopy (CryoEM) revealed a localized interface between OT8 and host cells, including tubular structures that may be type IV secretion system (T4SS) pili. These extend from the parasite toward the host envelope (Fig. 2e, Supplementary Fig. S5, Table S1 and Note 3), suggestive of direct access to host cellular material. As indirect evidence, infected Mc4 often exhibited shrunken cytoplasms, including detachment of the inner membrane from the outer sheath (Fig. 2f and Supplementary Fig. S6). Notably, at the OT8 attachment site, the host cytoplasmic membrane and outer sheath remained closely associated.

**Fig. 2.**
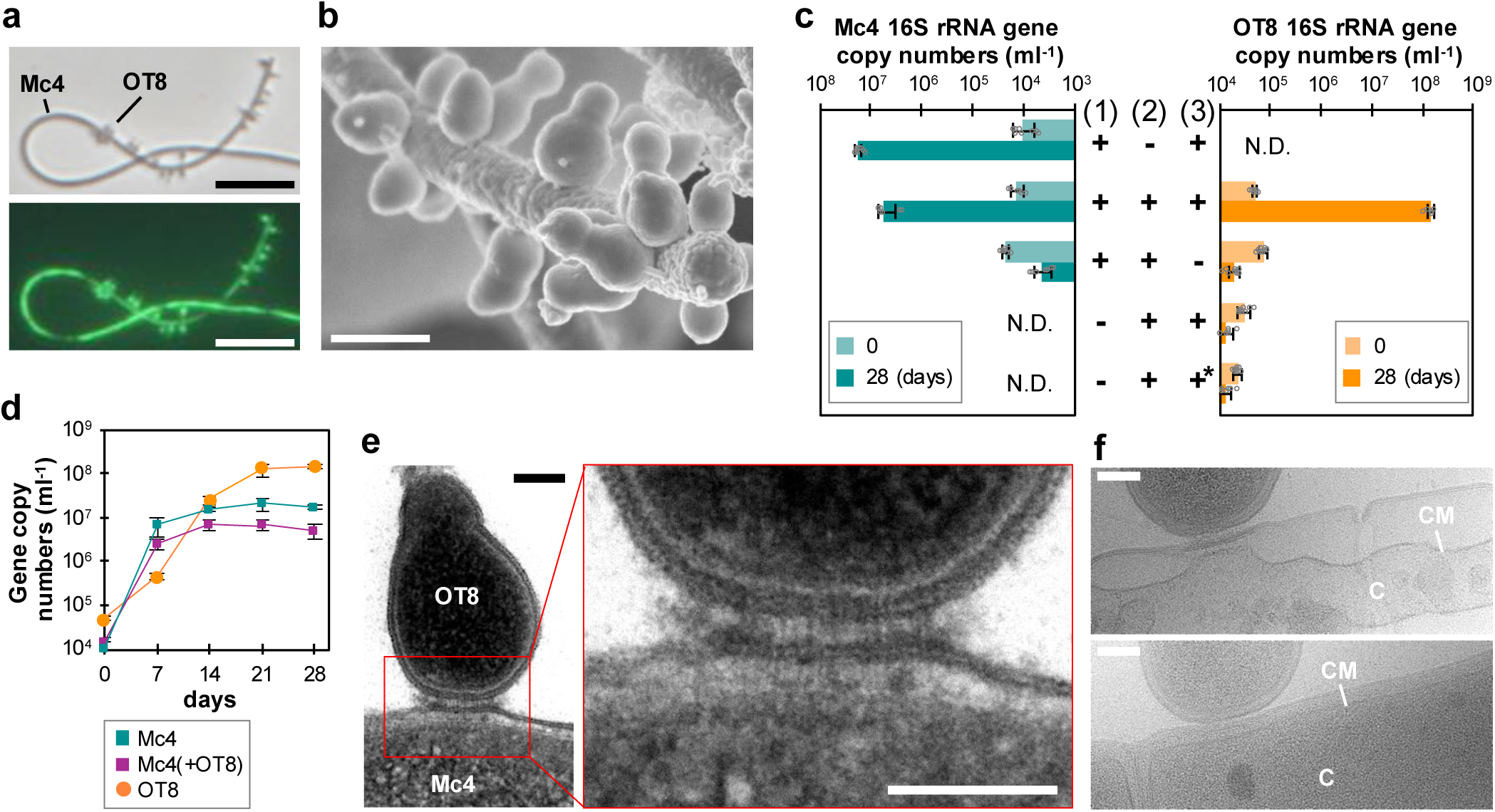
Parasitic interaction between strain OT8 and host strain Mc4. **a,b,** Cells imaged by phase-contrast microscopy (DNA staining in lower panel; **a**) and scanning electron microscopy (**b**). **c,d,** Growth in mono- and co-culture. Mc4 and OT8 abundances were quantified by 16S rRNA gene copy number across culture conditions (**c**) and over time (**d**). In **c**, columns (1), (2) and (3) indicate the presence (+) or absence (−) of Mc4 cells, OT8 cells and Mc4 growth substrates, respectively; asterisks mark cultures supplemented with Mc4 cell lysate instead of these substrates. **e,** Ultrathin-section transmission electron microscopic images showing tubular connections between OT8 and Mc4 cytoplasm. **f,** Cryo-electron microscopic images showing cytoplasmic shrinkage of parasitized Mc4 cells (top) compared with a normal cell (bottom). Scale bars, 5.0 μm (**a**); 400 mm (**b**); 100 nm (**e,f**).

To identify the host-derived material acquired by OT8, we examined changes in host macromolecules during infection. Mc4/OT8 co-cultures exhibited reduced host ribosomal RNA levels relative to uninfected Mc4 cultures, based on the ratio of Mc4 16S rRNA to 16S rRNA gene copy numbers (Fig. 3a). In parallel, incubation with the click-reactive guanosine analog AzG^17^, which fluoresces only when incorporated into RNA, produced signals in both Mc4 and attached OT8 cells, whereas host-free OT8 cells showed no detectable fluorescence (Fig. 3b–d and Supplementary Fig. S7). Because only Mc4 encodes the capacity to synthesize GTP from guanosine (Supplementary Fig. S8), these results indicate that OT8 acquires host-derived RNA during infection (Supplementary Note 4).

**Fig. 3.**
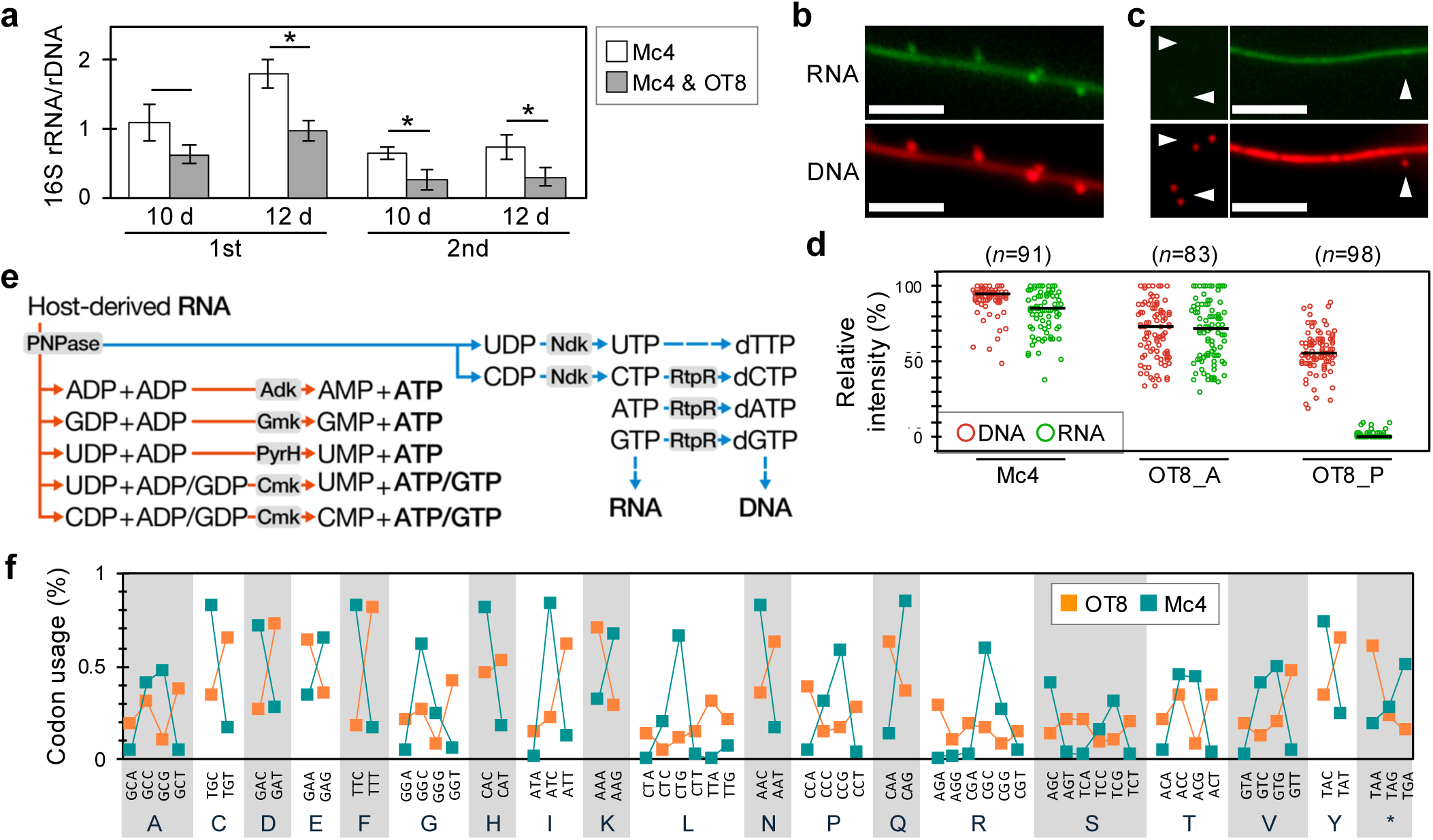
Acquisition and metabolic utilization of host-derived RNA by OT8. **a**, Ratio of Mc4 16S rRNA/16S rRNA gene (rDNA) copy number, used as an indicator of Mc4 rRNA abundance, in mono-cultures and co-cultures with OT8 at days 10 and 12 in two independent experiments. Bars and error bars indicate the mean and s.d. of technical replicates, respectively (*n*=3). Asterisks indicate *p* < 0.05 by two-sided Welch’s t-test. **b**-**d**, Visualization of host-derived RNA transfer to OT8 using a nucleoside analog AzG, which becomes fluoresces when incorporated into RNA polymers. Fluorescence shown for Mc4–OT8 co-cultures (**b**, top) and a mixture of Mc4 cells from an AzG-amended monoculture and host-detached OT8 cells (arrowheads) collected by filtration from an Mc4–OT8 co-culture and incubated with AzG (**c**, top). Fluorescence from SYTO59 targeting DNA are also shown (**b,c,** bottom). Scale bars, 5 μm. Background-subtracted fluorescence intensities of DNA and RNA signals in representative Mc4 cells, host-attached OT8 cells (OT8_A), and host-detached planktonic OT8 cells (OT8_P), normalized within each image to the brightest signal (**d**). Each point represents a single cell, and horizontal bars indicate medians. **e**, Proposed pathway for degradation of host-derived RNA leading to ATP/GTP production (orange) and RNA/DNA polymer synthesis (blue) in OT8. PNPase, polynucleotide phosphorylase; Adk, adenylate kinase; Gmk, guanylate kinase (absent from OT8); PyrH, uridylate kinase; Cmk, cytidylate kinase; Ndk, nucleoside-diphosphate kinase; RtpR, ribonucleoside-triphosphate reductase. **f**, Comparison of codon usage between OT8 and Mc4. Values indicate the relative usage frequency of each synonymous codon within the corresponding amino acid.

### Metabolic processing of RNA-derived nucleotides

Genomic reconstruction revealed that OT8 lacks pathways for *de novo* nucleotide biosynthesis (Supplementary Table S1) and is severely depleted in proteins associated with redox metabolism (Supplementary Table S2), canonical energy metabolism and catabolism of major host-derived major substrates, deoxyribonucleotides, amino acids, carbohydrates, and peptidoglycan-derived muropeptides (Supplementary Table S3). Instead, OT8 encodes phosphorylases, including polynucleotide phosphorylase (PNPase), and phosphotransferases such as adenylate kinase (Adk) and cytidylate kinase (Cmk), which together can cleave RNA and interconvert the liberated nucleotide species (Fig. 3e and Supplementary Table S3). These enzymes indicate that RNA-derived nucleotides are processed through intracellular metabolic reactions that redistribute phosphate groups among nucleotide pools, maintain nucleotide triphosphate levels, and provide entry points into RNA and DNA precursor biosynthesis (Fig. 3e). Cmk may play a particularly important role given its capacity to exploit all four NDP types^18^ (Fig. 3e). Consistent with the physiological importance of this pathway, PNPase and Cmk were among the top 5% of genes optimized for expression in OT8 (Supplementary Table S3).

In this context, host-derived RNA may support cellular energetics by supplying pre-energized phosphate-bearing metabolites that can be redistributed through intracellular nucleotide interconversion reactions. Although direct measurements of energy flux are not possible in this system, the combination of RNA uptake capacity, intracellular nucleotide-processing capacity, and the absence of conventional energy metabolism pathways supports a model in which host-derived RNA contribute to cellular energetics.

Energetic estimations further indicated that host-derived RNA—either total RNA or ribosomal RNA—could satisfy both the energetic and biosynthetic requirements for OT8 cell replication (Supplementary Note 5). We estimate that the total RNA and rRNA content of a single Mc4 cell could each support division of an OT8 cell, based on cell size and biosynthetic pathway constraints (see Methods). Moreover, assuming continuous RNA synthesis and consumption rates consistent with the doubling times and RNA contents of Mc4 and OT8, a single Mc4 cell could theoretically sustain replication of at least four OT8 cells. In contrast, the intracellular pools and synthesis rates of tRNA, mRNA, and free nucleotides alone were insufficient to satisfy the predicted energetic and biosynthetic demands of OT8 replication.

The resulting RNA-catabolic route represents a highly streamlined strategy, allowing NTP regeneration from RNA-derived nucleotide intermediates through as few as two enzymatic steps (Fig. 3e). Unlike several minimal energy-conserving metabolisms (*e.g.*, pyruvate disproportionation or carboxylate decarboxylation), this route avoids synthesis of expensive organic cofactors. This metabolic strategy effectively externalizes canonical redox-coupled energy conservation to the host by directly exploiting the chemical energy stored in host-derived RNA.

This metabolic strategy is also distinct from well-known cell-invading parasites that assimilate host nucleotides. Unlike OT8, these organisms’ membranes directly contact the host cytosol and can thus use transporters to uptake host nucleotides (which are absent in *Minisyncoccota*/Patescibacteriota). Further, these exclusively parasitize eukaryotic cells in which nucleotide pools are presumably orders larger than bacterial cells and better suited to sustain parasite cell division. Notably, host-attached OT8 produced progeny even in the absence of host growth substrates where the host is unlikely to be synthesizing nucleotides *de novo* (Supplementary Fig. S3), further supporting that OT8 likely employs host RNA rather than host NTPs as a nucleotide source.

### Codon utilization is complementary between the parasite and host

Further comparing the genomes of OT8 and Mc4, we found complementary codon usage patterns (Fig. 3f) despite the close phylogenetic relationship between *Chloroflexota* and *Minisyncoccota*/Patescibacteriota^1,19,20^. If Mc4-derived mRNAs were assimilated and translated by OT8, these mRNAs would exhibit markedly low expression potentials, based on gene expressivity analysis^21^ (Supplementary Fig. S9). Organisms and their genes are generally optimized to specific codon usage and deviation from this can negatively influence translation efficiency and accuracy^22,23^. Thus, this difference in codon utilization would theoretically minimize risk of OT8 translation of Mc4-derived mRNA.

### Conservation of nucleotide-centered metabolism across the phylum

To determine whether the metabolic features observed in OT8 are broadly conserved across the phylum, we compared gene inventories across diverse representatives of *Minisyncoccota*/Patescibacteriota. This analysis revealed recurrent depletion of both redox-associated proteins and *de novo* nucleotide biosynthesis pathways, coupled with retention of enzymes inferred to support RNA metabolism in OT8. Most genomes lack pathways for *de novo* nucleotide biosynthesis [93% of genomes examined, Fig. 4a(1)]. Nevertheless, 78% of genomes harbor genes encoding enzymes that could facilitate NTP generation from RNA—PNPase and Adk [Fig. 4a(2)]. Separately, 51% of genomes harbors both PNPase and Cmk that would theoretically allow utilization of all RNA-derived NDPs [Fig. 4a(3)]. This includes well-studied species such as *Ca*. Absconditicoccus praedator^24^, *Ca*. Vampirococcus lugosii^25^ and *Ca*. Southlakia epibionticum^26^. Like OT8, these species had distinct codon usage patterns from their respective hosts and exhibited low expression potential for host genes (Supplementary Fig. S9 and S10). Notably, the magnitude of this difference is smaller for parasite-host pairs in which the parasite lacks the genomic potential for RNA catabolism (Supplementary Fig. S9). Together, these patterns indicate a widespread capacity to utilize exogenous RNA as a metabolic resource.

**Fig. 4.**
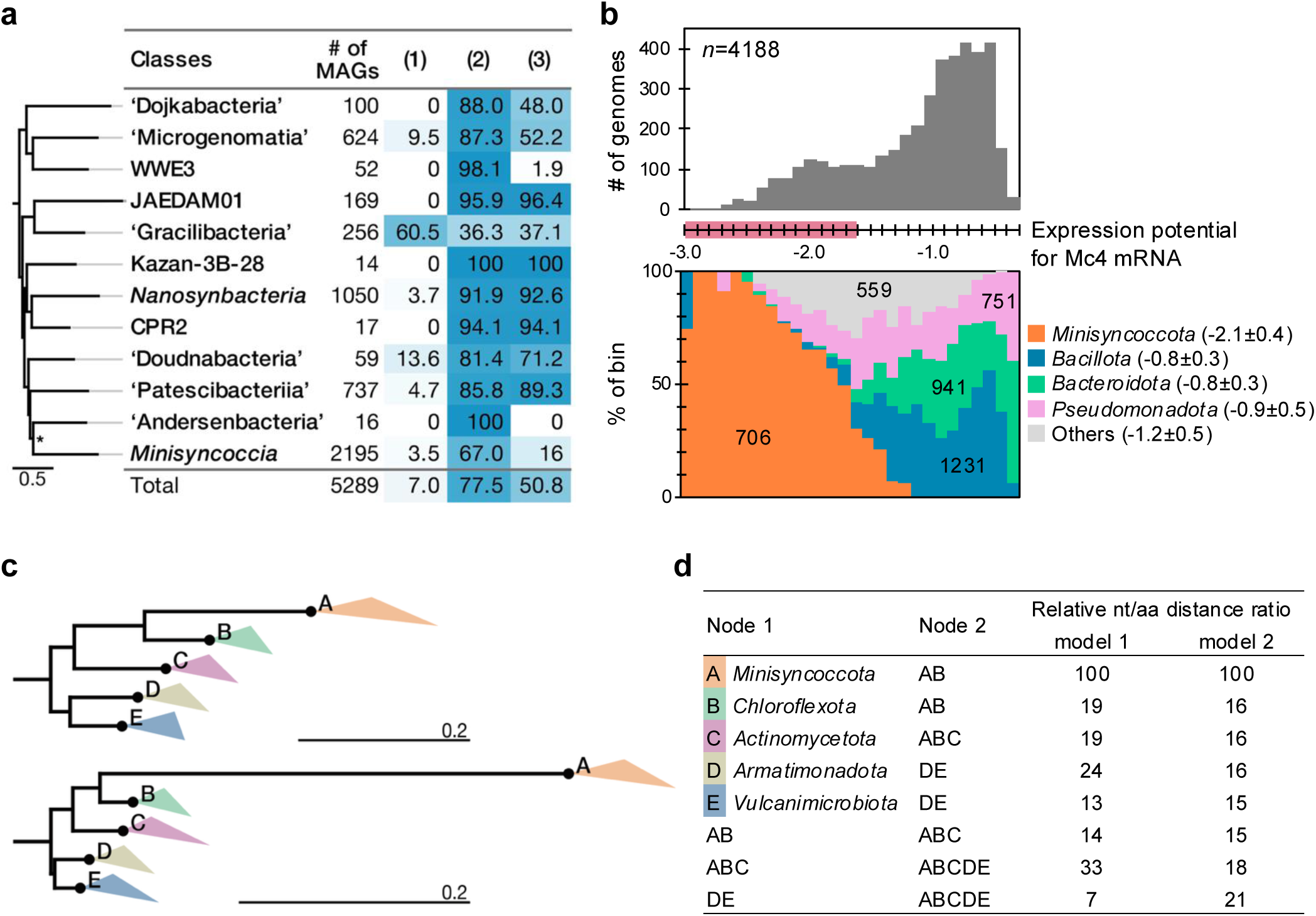
Evolutionary signatures of RNA-dependent nucleotide metabolism and codon-usage complementarity in *Minisyncoccota*/Patescibacteriota. **a,** Distribution of nucleotide metabolic capacities across major *Minisyncoccota*/Patescibacteriota classes: *de novo* nucleotide biosynthesis (1), potential coupling of RNA degradation to NTP generation through PNPase and Adk (2) or PNPase and Cmk (3). Values indicate the percentage of genomes in each class encoding the corresponding capacity. **b,** Distribution of expression potential for Mc4 mRNAs evaluated against bacteria with genomic G+C contents similar to OT8 (38.8–42.8%). The red range in x-axis corresponds to the range of expression potential observed for host-parasite pairs with predicted RNA-catabolic capacity (Supplementary Fig. S9). The lower panel shows the taxonomic composition of genomes in each bin. Numbers indicate genome counts for each lineage, and values in parentheses indicate median±SD expression potential. **c,** Maximum-likelihood phylogenies of *Bacillati* phyla reconstructed from a concatenated alignment of conserved marker proteins using amino-acid sequences (upper) and the corresponding nucleotide sequences (lower). Black circles indicate the last common ancestor of each collapsed phylum. The scale bar applies to the black backbone branches only. Colored triangles are horizontally compressed for visualization. **d**, Ratios of nucleotide (nt)- to amino-acid (aa)-based evolutionary distances between *Bacillati* lineages and their internal ancestral nodes, calculated from the trees in **c**. Distances are reported as relative values normalized such that the distance between node A and node AB is set to 100. Values were calculated under two different models: LG+60+C+F (model 1) and Poisson+UDM0064LCLR (model 2).

### RNA metabolism may be an ancestral feature

*Minisyncoccota*/Patescibacteriota have AT-rich genomes (median G+C content of 42.1%), whereas Mc4 and other previously identified hosts have GC-rich genomes, raising the possibility that codon-usage complementarity reflects differences in G+C contents. However, comparison of OT8 and other *Minisyncoccota* with bacterial taxa with similar G+C contents (39.3–42.3%) showed that *Minisyncoccota* retain distinct codon-usage profiles in ribosomal protein genes (Supplementary Fig. S11 and S12) and lower codon-usage compatibility with genes from Mc4 and other previously identified host than did other taxa, based on expression-potential estimates^21^ (Fig. 4b and Supplementary Figs. S13). Meanwhile, these *Minisyncoccota* also broadly showed low expression potential for genes from bacteria with Mc4-like G+C contents (63.4–67.4%) (Supplementary Fig. S14). Together, these analyses indicate that the codon-usage complementarity of *Minisyncoccota* is both lineage-specific and broadly observed across phylogenetically diverse GC-rich genomes, rather than simply reflecting genome-wide G+C content differences (Supplementary Note 6). This broad complementarity suggests that RNA-metabolizing *Minisyncoccota* are pre-adapted to interact with GC-rich hosts, potentially reducing constraints on host switching during evolution.

*Minisyncoccota*/Patescibacteriota generally possess AT-rich genomes yet phylogenetically nest within a clade of GC-rich lineages, including *Chloroflexota*, *Actinomycetota*, *Vulcanimicrobiota*, and *Armatimonadota*^1,19,20^ (median G+C contents of 56.7%, 65.6%, 62.9%, and 59.6% respectively). This contrast suggests a sharp decrease in G+C content along the evolutional trajectory leading to *Minisyncoccota*. Phylogenetic analyses further revealed an elevated rate of nucleotide substitutions relative to amino acid substitutions along this branch (Fig. 4c, d), suggesting selection favoring silent mutations substituting G/C with A/T nucleotides. Because most members of this phylum generally lack *de nov*o nucleotide biosynthesis pathways, and nucleotide biosynthesis genes show scattered phylogenetic distributions (Supplementary Fig. S15), this shift is unlikely to reflect reduced biosynthetic cost^27,28^. Instead, these observations suggest that increasing AT richness may have been driven by for codon complementation in the context of parasitism-mediated exposure to host-derived mRNA. Consistent with this model, *Minisyncoccota* formed monophyletic clades in phylogenies of PNPase and VirB4 (Supplementary Fig. S16), each presumably associated with RNA catabolism and macromolecule transfer, suggesting ancient acquisition of these functions.

Together, these suggest that RNA-based parasitism may be an ancestral feature of this lineage. The observations warrant revisiting the unexplained prevalence of unusual features in the information-processing machinery of *Minisyncoccota*/Patescibacteriota (*e.g.*, introns in ribosomal RNA) ^4,6,29–32^ as systems that may support the utilization of exogenous RNA.

### A putative model for host RNA uptake

The mechanism by which OT8 and related organisms acquire RNA from their hosts remains unresolved. Based on cryoEM, OT8’s tubular structures appear to penetrate the host envelope (Fig. 2e). Together with host cytoplasmic shrinkage observed for infected host cells (Fig. 2f), these findings suggest that OT8 may acquire host cytoplasmic material, including RNA, through direct physical interaction with its host.

This interpretation is further supported by the growth morphology of host-attached OT8 cells, which enlarge upon attachment and appear to use the increased cell volume to produce daughter cells (Supplementary Fig. S17 and Note 7). These observations suggest that host-derived material is transferred during parasite cell enlargement and growth, though the underlying mechanism remains to be determined. Notably, we found that most members of this phylum and known hosts (including OT8 and Mc4 respectively) lack discernible aquaporins (Supplementary Fig. S18), indicating that restricted water transport^33^ may play a role.

It is important to note that other transport routes for host cellular material may exist. For lipid synthesis, OT8 lacks the genetic capacity for *de novo* phospholipid synthesis, including those universally conserved across other bacterial phyla (Supplementary Table S1). Analyzing the fatty acid compositions extracted from cells in axenic Mc4 cultures and Mc4-OT8 co-cultures, we found that the fatty acid profiles were nearly identical (Supplementary Fig. S19), suggesting that OT8 directly utilizes lipid molecules synthesized by its host. Unlike other cellular components sourced from the host, such as RNA, phospholipids are only present in the cellular membrane and cannot be assimilated from the cytoplasm. These observations suggest that lipid transfer occurs through a mechanism distinct from RNA acquisition.

### Conclusion

Here we describe an organism that reveals a novel extreme in the broad spectrum of life’s strategies: the reduction of core metabolic functions—canonical electron-transfer-based energy metabolism, broader redox metabolism, and *de novo* nucleotide synthesis—through exploitation of another organism’s RNA as a metabolic resource. This strategy arises from a cellular and genomic architecture specialized for acquiring and degrading host-derived RNA, and genomic signatures of this metabolism are widely conserved and likely ancestral across *Minisyncoccota*/Patescibacteriota. By reorganizing cellular metabolism around a host-derived nucleic acid polymer, this lineage expands the known strategies of parasitism to include direct metabolic exploitation of informational molecules. This provides critical insight into the physiology underlying the evolutionary success of these metabolically reduced organisms and distinctive features of their information-processing systems^4,6,29–32^. More broadly, these findings demonstrate that cellular metabolism can be sustained through direct exploitation of informational macromolecules, expanding the known principles by which life acquires energy and builds biomass.

## Materials and Methods

### Genomic analyses

Prokaryotic genomes were obtained from GTDB release 10-RS226^34^. We retained genomes designated as GTDB representatives and further filtered them using genome quality criteria of ≥85% completeness and ≤5% contamination. Coding sequences were identified using prodigal v2.6.3^35^ and functionally annotated using eggNOG-mapper v2.1.12 against the eggnog DB v5.0.2^36^ using default settings. The resulting functional annotations, including KEGG Orthology identifiers and Pfam domain assignments, were used for downstream protein-count analyses. To estimate the genomic capacity for nucleotide biosynthesis, we counted proteins associated with *de novo* purine and pyrimidine biosynthesis pathways based on KEGG module definitions. Specifically, KEGG Orthology identifiers assigned by eggNOG-mapper were compared against the KO sets included in KEGG modules M00048 and M00051, corresponding to purine biosynthesis and pyrimidine biosynthesis, respectively. For each genome, proteins annotated with KOs present in either of these modules were counted as nucleotide biosynthesis proteins. Redox-dependent proteins were identified using a curated Pfam-domain-based screening strategy. Briefly, Pfam domains associated with experimentally characterized redox-related UniProtKB entries were compiled into a redox-associated Pfam marker set. The complete list of Pfam domains, domain combinations, and UniProtKB evidence entries used for this analysis is provided in Supplementary Table S4. Pfam annotations assigned by eggNOG-mapper were searched against this marker set, and proteins containing matching redox-associated Pfam domains were counted as putative redox-dependent proteins. For marker patterns consisting of multiple Pfam domains, all listed domains were required to be present in the same protein. Distinct marker patterns were treated as alternative markers. Energy metabolism-associated proteins were counted using an analogous KEGG module-based approach. KEGG Orthology identifiers assigned by eggNOG-mapper were compared against KO sets from KEGG modules associated with canonical energy metabolism pathways, including M00144, M00151, M00152, M00153, M00155, M00156, M00174, M00356, M00357, M00416, M00417, M00528, M00529, M00530, M00563, M00567, M00596, M00804, M00973, M00984, M00985, and M00986. Proteins annotated with KOs present in these modules were counted as energy metabolism-associated proteins for each genome. ATP/ADP translocase-like nucleotide transport systems were identified based on Pfam domain assignments from the eggNOG-mapper annotations. Specifically, proteins containing Pfam domains associated with ATP/ADP translocases, including PF08449, PF00153, and PF03219, were searched in the Pfam annotations assigned to each genome-encoded protein.

Genomic DNA was extracted from the pure culture of Mc4 and co-culture of Mc4 and OT8 using the Blood and Cell Culture DNA Maxi Kit (QIAGEN, Netherlands) according to the manufacturer’s instructions. DNA was sequenced using a PacBio Revio system (Pacific Biosciences, USA) and DNBSEQ G-400 system (MGI Tech Co., Ltd., China) at Bioengineering Lab. Co., Ltd. Hybrid sequence assembly was carried out using Unicycler v0.5.1^37,38^.

For genomic sequences of OT8 and Mc4, coding sequences were identified using prodigal and functionally annotated using eggNOG-mapper, as described above, and CD-search against the CDD v3.21 database^39^ using E-value threshold of 1×10^-5^ and “Standard” result mode. For CAZy annotation, carbohydrate-active enzymes (CAZymes) were identified with DRAM (v1.5.0) using default settings (HMM-based searches against CAZy-associated profiles). For Supplementary Table S3, we report presence/absence of functional categories (defined as ≥1 CAZy hit assigned to a given category) rather than listing all individual CAZy genes.

A set of conserved marker proteins involved in replication, transcription, and translation representing a subset of the GTDB marker protein set^40^ was annotated for OT8, Mc4, and related species, as well as metagenome-assembled genomes (MAGs) with ≥85% completeness and ≤5% contamination. Annotation was performed using MMseqs2 v17.b804r^41^ in “search” mode (--start-sens 4 -s 8.5 --sens-steps 3 --cov-mode 2 --max-seqs 50000 -c 0.7 --min-seq-id 0.25 --filter-hits) against the GTDB marker protein database. The identified sequences were aligned using MAFFT v7.49^42^ using the L-INS-i mode (--localpair --maxiterate 500), concatenated, and trimmed with BMGE v1.12^43^ using the parameters -g 0.67 -m BLOSUM30 -b 3. A maximum likelihood tree was inferred with IQ-TREE v2.2.6^44^ using the LG+C60+G+F model, with branch support assessed using 1,000 ultrafast bootstrap replicates and 1,000 SH-like approximate likelihood ratio test replicates (SH-aLRT). For phylogeny of PNPase, and VirB4 (conserved ATPase of T4SS) and major intrinsic proteins, including aquaporins, tree was constructed with IQ-TREE employing the LG+F+R10+PMSF model and 1,000 ultrafast bootstrap and SH-aLRT replicates. For amino acid- and nucleotid-based tree comparison, the aligned and trimmed amino acid positions were applied to the corresponding nucleotide sequences. For amino acid-based tree construction, the Poisson+UDM0064LCLR model was also used. For nucleotide-based tree construction, the GTR+F+I+R7 model, selected via ModelFinder, was employed under the constraint of the amino acid-based tree topology using IQ-TREE. Amino acid sequences of proteins, adenylosuccinate synthase, adenylosuccinate lyase, amidophosphoribosyltransferase, aspartate carbamoyltransferase, dihydroorotase, and orotidine-5’-phosphate decarboxylase, used as markers of *de novo* purine and pyrimidine biosynthesis, were aligned using MAFFT and trimmed by trimAl v1.5^45^. The maximum likelihood tree was constructed based on these protein sequences using FastTree ver.2.2.0^46^.

Codon usage–based gene expressivity was evaluated using the Measure Independent of Length and Composition (MILC) and MILC-based Expression Level Predictor (MELP) metrics^21^. MELP scores of coding sequences predicted from the OT8 genome was calculated using the R package coRdon v1.16.0^47^ based on the codon usage pattern of ribosomal protein genes as the highly expressed reference set. Genes were then ranked by MELP score, and the percentile rank (“optimized genes”) for each target gene (listed in Supplementary Table S3) was calculated. The averaged codon usage pattern in all coding-sequences in a genome was calculated using a CUSP program in the EMBOSS package v.6.6.0^48^. For comparison of codon utilization between *Minisyncoccota*/Patescibacteriota and their hosts, the host genome sequence of *Ca*. V. lugosii is unavailable. Accordingly, a closely related species, *Halorhodospira halophila*, identified from 16S rRNA gene similarity^25^ was used. To estimate the potential expressivity of host-derived mRNA within parasitic *Minisyncoccota* cells (*i.e.*, MELP values of host genes under the *Minisyncoccota* translational context), the ribosomal protein genes from *Minisyncoccota* genomes were used as the reference set, whereas the corresponding host genes were included in the calculation as target genes but not designated as reference sequences. For analyses shown in Supplementary Fig. S14, we evaluated host-like (Mc4-like) G+C genomes (63.4–67.4%) under three *Minisyncoccota* translational backgrounds using ribosomal protein reference sets derived from OT8, *Minisyncoccia* (GCA_021734905; 41.1% G+C), and *Ca*. Dojkabacteria (GCA_016932785; 40.5% G+C) Prior to MELP calculation, coding sequences shorter than 240 bp and sequences containing ambiguous nucleotides (N) were excluded because MELP is not computed for such genes in the employed pipeline. In addition, MELP computation requires that the reference set covers all codons used by target genes; therefore, for each *Minisyncoccota* genome, we ensured complete codon coverage in the reference by appending the missing codon(s) (as synonymous triplets) to a single ribosomal protein gene sequence in the reference set. This codon-completion step did not alter the overall genome-wide distribution and ranking trends of gene expressivity scores, as confirmed by recalculating MELP without codon completion for some genomes.

For comparison of ribosomal protein codon usage, relative synonymous codon usage (RSCU) was aggregated using only ribosomal protein subunits present in ≥97% of genomes (33 subunits), and only genomes encoding at least 30 of these 33 subunits were included to minimize biases due to incomplete recovery of ribosomal gene. Within-class correspondence analysis (WCA) was then performed on the resulting ribosomal protein RSCU profiles. For multivariate comparisons between *Minisyncoccota* and non-*Minisyncoccota* genomes, coordinates on WCA axes 1–3, which together explained 61.6% of the variance, were used for PERMANOVA and PERMDISP analyses.

The presence or absence of *de novo* nucleotide synthesis in *Minisyncoccota* members was determined using the functional annotation dataset generated by eggNOG-mapper (as described above), which was assessed against the KEGG modules for purine and pyrimidine biosynthesis (M00005, M00048–M00052). Because the analyzed genomes were not always complete, a pathway was considered present when a genome encoded all but at most one of the required enzymes (*i.e.*, allowing one missing essential gene). Genomes meeting this criterion were classified as possessing *de novo* nucleotide biosynthesis [as shown in Fig. 4a(1)]. Genomes lacking *de novo* nucleotide biosynthesis were further screened for additional metabolic capacities based on the presence of marker genes in the annotation dataset: RNA-derived NTP generation was assigned to genomes encoding PNPase and Adk [Fig. 4a(2)] or those encoding PNPase and Cmk [Fig. 4a(3)]. These genes were identified in the functional annotation datasets generated by CD-search, as described above.

### Environmental sample collection and cultivation

The sediment and formation water samples were collected from a settling pond that was placed downstream of a commercial natural gas and water producing well to remove suspended sand particles from the formation water in Mobara, Chiba prefecture, Japan^49^. The samples came from the gas-bearing aquifers that consist of repeating sequences of turbidite (alternating beds of sandstone and mudstone). These sediments were deposited in deep marine environments during the Plio-Pleistocene periods^45^. The water temperature was 19.1°C, the pH was 7.5, and the redox potential was -248 mV. The Cl^-^concentration was 13,000 mg l^-1^, and the sulfate concentration was 82 mg l^-1^. The natural gases produced in this area composed mainly of methane (99%), and the origin of methane was suggested to be of biogenic based on stable isotopic analysis^45^. The sample collection was performed by authors with permission from the Kanto Natural Gas Development Co., Ltd.

The samples were collected in sterilized glass bottles with butyl rubber stoppers and screw caps. The bottle was purged with N_2_ gas prior to sample collection and was filled with the water to maintain the samples under anaerobic conditions. The sediment and formation water samples were collected at a 1:2 volume ratio.

Anaerobic enrichment cultivation of sediment and formation water samples was conducted as previously described^15^, with modifications. Sediment samples were mixed with formation water to prepare slurry samples within an anaerobic chamber. Aliquots of 20 ml of the slurry were dispensed into 70-ml serum vials, sealed with butyl rubber stoppers and aluminum crimps, all under anaerobic conditions. The vials were incubated without the addition of exogenous nutrients under an N_2_/CO_2_ (80:20) gas atmosphere at 25°C. After approximately 1.5 years of incubation, 2 ml of the methane-producing culture from the slurry was inoculated into a saline mineral medium^50^ supplemented with 0.1 mM titanium(III) citrate as a reducing agent. Cultivation was carried out in 75-ml serum vials containing 20 ml of medium, incubated at 45°C under the same N_2_/CO_2_ (80:20) atmosphere. The enrichment cultures underwent two successive transfers at 4-month intervals. From one of the duplicate cultures, microbial cells were harvested by centrifugation, resuspended in 4 ml of fresh saline mineral medium in a serum vial pre-filled with N_2_ gas, and autoclaved at 121°C for 10 minutes. The resulting sterilized biomass was then used as a nutrient source for the corresponding duplicate enrichment culture. This subculturing process was repeated six times, each at 4-month intervals.

From the final passage culture, individual cells targeting a filamentous bacterium affiliated with the family *Aggregatilineaceae* were isolated in pure culture using a deep agar slant method [8 g l⁻¹ noble agar (BD Difco, USA) supplemented with autoclaved biomass from the parent culture] in combination with the dilution-to-extinction method^50^. Following three months of incubation, a single colony was transferred to fresh liquid medium. Despite this isolation strategy, the filamentous *Aggregatilineaceae* bacterium persisted in co-culture with a morphologically distinct irregular-rod-shaped bacterium assigned to the family *Thermovirgaceae* (phylum *Synagistota*). To obtain a pure culture of the *Thermovirgaceae* strain, three successive rounds of isolation using the deep agar slant method were performed. The resulting isolate was designated strain Ths. A cell-free culture supernatant of Ths together with 0.5 g l⁻¹ yeast extract (BD Difco) was subsequently added to the co-culture and subjected to additional rounds of purification using the same method. After three iterations, a pure culture of the *Aggregatilineaceae* strain, designated Mc4, was obtained. Purity of Mc4 culture was confirmed by phase-contrast microscopy and the absence of contaminating sequences in whole-genome DNA sequencing data, as described below.

Strain Mc4 was cultivated at 40°C in saline mineral medium supplemented with 1.0 g l⁻¹ yeast extract, Ths culture supernatant (4 ml of this supplemented medium was added to 18 ml of base medium) and 0.1 mM titanium(III) citrate (as a reducing agent). Strain Ths was cultivated at 40°C in saline mineral medium containing 1.0 g l⁻¹ yeast extract, 2.0 g l⁻¹ tryptone (BD Difco), and 0.1 mM titanium(III) citrate. After 7-days cultivation, cell-free culture supernatant was obtained by centrifugation followed by sterilization through a 0.2-μm pore-size filter.

Under light microscopy, ultrasmall cells were observed attached to Mc4-like filamentous bacteria in the final passage culture used for the attempted isolation of strain Mc4, as described above. To establish a co-culture of the ultrasmall cells and Mc4, this final passage culture was sequentially filtered through 0.4-μm and 0.2-μm pore-size filters. The filtrate was then inoculated into a pure culture of Mc4. After three successive passages using this procedure, a stable co-culture was obtained, consisting of Mc4 and ultrasmall cells belonging to the candidate class *Ca*. Microgenomatia, designated as strain OT8. Purity of this co-culture was confirmed by phase-contrast microscopy and the absence of contaminating sequences in 16S rRNA gene amplicon and whole-genome DNA sequencing data, as described below. The cultivation medium and conditions for the co-culture of OT8 and Mc4 were identical to those used for the pure culture of Mc4, as described above.

### Amplicon sequencing analysis

Total DNA was extracted from enrichment cultures and isolated strains using the ISOSPIN Fecal DNA Kit (NIPPON GENE, Japan), following the manufacturer’s instructions. The V4 region of the 16S rRNA gene was amplified using the primer pair

515F and 806R, and sequenced on the Illumina MiSeq platform at Bioengineering Lab. Co., Ltd., Japan. The resulting sequence reads were processed using QIIME2 version 2024.10^51^ under standard protocols and taxonomically classified using the SILVA rRNA gene database release 138.2^52^.

### Microscopy

Cell morphology and ultrastructure were analyzed using phase-contrast and fluorescence microscopy (BX51; Olympus, Japan), confocal laser scanning microscopy (LSM800; ZEISS, Germany), scanning electron microscopy (SEM) (JSM-7500F; JEOL, Japan), cryo-electron microscopy (CryoEM) (CRYO ARM 300 II; JEOL), and transmission electron microscopy (TEM) [H-7600; Hitachi, Japan, for negative staining; JEM-1400Plus; JEOL, for ultrathin sectioning].

For fluorescence microscopy, genomic DNA was stained with SYBR Green I (Thermo Fisher Scientific, USA) at a final concentration of 2 µg ml⁻¹ and visualized using fluorescence microscopy.

Peptidoglycan was labeled with the fluorescent D-amino acid HADA (Tocris Bioscience, UK). Mc4 and OT8 cells, previously grown separately in pure and co-cultures for 7 and 16 days, respectively, were inoculated into fresh medium supplemented with 1 mM HADA and incubated at 40°C for 2.5 days. The cells were then harvested and washed twice with PBS. Genomic DNA was subsequently stained with SYTO59 (Thermo Fisher Scientific) at a final concentration of 5 µM and incubated for 20 minutes at room temperature. Samples were imaged by confocal laser scanning microscopy and analyzed by fluorescence microscopy for cell counting.

For RNA labeling, three experimental conditions were prepared: Mc4–OT8 co-cultures, Mc4 monocultures, and host-free OT8 cell preparations. Host-free OT8 cells were obtained from an 18-day co-culture of Mc4 and OT8 by filtration through 0.4-μm pore-size filters. Each preparation was incubated in medium containing 250 µM 2’-deoxy-2’-azidoguanosine (AzG; Angene International Ltd., China) at 40 °C for 3 days. After incubation, cells were harvested, washed with PBS, and fixed with 4% paraformaldehyde (PFA) for 15 minutes at room temperature. Following two PBS washes, cells were permeabilized with 0.1% Triton X-100 for 15 minutes and washed again twice with PBS. For the control preparation shown in Fig. 2c, AzG-incubated Mc4 monoculture cells and AzG-incubated host-free OT8 cells were mixed before click labeling. Mc4 monoculture cells were included in this control mixture to verify that the click-labeling reaction functioned in the same preparation. Click labeling was then performed using Alexa Fluor 488 alkyne in the presence of a copper catalyst, according to the manufacturer’s protocol (Click-iT Cell Reaction Buffer Kit; Thermo Fisher Scientific) for 30 minutes at room temperature. After three PBS washes, DNA was stained with SYTO59 and visualized by fluorescence microscopy. Three biological replicates were observed (∼30 cells per replicate). For quantification, background-subtracted mean fluorescence intensity was measured for individual cells. Relative fluorescence intensity (%) was calculated by normalizing each cell’s intensity to the brightest labeled cell in the same image (either Mc4 or OT8), which was set to 100.

For SEM, cells were prefixed with 2% glutaraldehyde in 0.1 M sodium cacodylate buffer (pH 7.4) at 4 °C overnight. Post-fixation was performed sequentially with 1% tannic acid for 2 h at 4 °C and 2% osmium tetroxide for 2 h at 4 °C. The samples were dehydrated through a graded ethanol series, lyophilized, sputter-coated with osmium and observed at Tokai Electron Microscopy, Inc., Japan.

For CryoEM, a 2 µl aliquot of concentrated co-cultured cells (OT8 and Mc4) was applied onto glow-discharged holey carbon grids (Quantifoil R 1/4 Cu; Quantifoil MicroTools GmbH, Germany). Grids were automatically blotted at 22 °C and 80% humidity and vitrified by plunging into liquid ethane using a Leica EM GP2 (Leica, Austria). Cryo-grids were mounted on a dedicated cartridge and loaded into a CRYO ARM 300 II electron microscope equipped with a cold field-emission gun operated at 300 kV, a hole-free phase plate^53^, and an in-column omega-type energy filter (30 eV slit width). Images were acquired on a K3 direct detection camera (GATAN, USA) at nominal magnifications of 10,000× to 15,000×, corresponding to a resolution of 3.3–4.9 Å per pixel, with a total electron dose below 1.5 e⁻/Å² using a low-dose system. Acquisition was performed using Minimum Dose System v4.2.12.34955.

For TEM, cells were subjected to freeze substitution using a solution containing 2% glutaraldehyde, 1% tannic acid in ethanol, and 2% distilled water at −80°C for 2 days with liquefied propane as the refrigerant. Following substitution, the cells were dehydrated overnight in absolute ethanol. For resin infiltration and embedding, the samples were treated twice with propylene oxide at room temperature for 30 minutes, followed by incubation in a 7:3 mixture of propylene oxide and Quetol-812 resin (Nisshin EM Co., Japan) at room temperature for 1 hour. The propylene oxide was then evaporated by overnight exposure at room temperature. Ultrathin sections were prepared using an ultramicrotome (Ultracut UCT, Leica), mounted on copper grids, stained with 2% uranyl acetate for 15 minutes and lead stain solution (Sigma-Aldrich, USA) for 3 minutes, both at room temperature, and observed at Tokai Electron Microscopy, Inc.

For negative staining, cells were fixed with 2.5% glutaraldehyde in 0.1 M sodium cacodylate buffer (pH 7.4) at 4 °C for 3 h, followed by post-fixation with 1% osmium tetroxide for 90 min at 4 °C. The fixed cells were applied onto carbon-coated grids, stained with 0.5% aqueous uranyl acetate at room temperature for 1 h, air-dried, and observed under TEM.

### Physiological characterization

All physiological experiments were conducted in triplicate. The growth of Mc4 and OT8 was monitored by quantifying 16S rRNA gene copy numbers using quantitative PCR, as described below. For morphological and structural observations, Mc4 and OT8 cells were harvested from a co-culture incubated for 21 days. To evaluate the growth behavior of Mc4 and OT8 (corresponding to the results shown in Fig. 2c), cultures were incubated either in the presence or absence of Mc4-specific growth substrates (as described above) or in the presence of Mc4 cell extracts. To prepare the cell extracts, Mc4 cells were harvested from 10 ml of pure culture, resuspended in 1 ml of saline mineral medium, and disrupted using a FastPrep-24 Classic Grinder (MP Biomedicals, USA) with three 60-second cycles at 5.0 m/s. For the inoculation of Mc4-free OT8 cells, co-cultures of Mc4 and OT8 were filtered through 0.2-μm pore-size filters.

To investigate the growth status of Mc4 during OT8 parasitism and growth (corresponding to the results shown in Supplementary Fig. S3), Mc4 and OT8 cells were harvested from a co-culture incubated for 1 week, washed twice with 10 mM phosphate-buffered saline (PBS, pH 7.2), and inoculated into fresh saline mineral medium either in the presence or absence of washed Mc4 cells (obtained from a 1-week pure culture) or Mc4 growth substrates.

To assess the dependency of OT8 on Mc4 (corresponding to the results shown in Supplementary Fig. S4), Mc4-free OT8 cells were obtained from an 18-day co-culture by filtration through 0.4-μm pore-size filters and inoculated into saline mineral medium supplemented with Mc4 growth substrates. These cultures were incubated at 40 °C for 0, 2, or 4 weeks, after which they were transferred to saline mineral medium containing both Mc4 cells and Mc4 growth substrates.

To assess the host range of OT8, we tested whether it could parasitize *Aggregatilinea lenta* strain MO-CFX2, the closest cultivated relative of Mc4. MO-CFX2 (=JCM 32065), obtained from the Japan Collection of Microorganisms (RIKEN BioResource Center, Japan) was grown under the same anaerobic conditions and medium composition as those used for Mc4, except that cultures were supplemented with substrates supporting MO-CFX2 growth, i.e., 0.1% yeast extract, 0.1% tryptone and 10 mM pyruvate^54^. Exponentially growing MO-CFX2 cells were incubated with Mc4-free OT8 cells at 30 °C under an N₂/CO₂ (80:20) atmosphere.

### Real-time PCR

To monitor the growth of Mc4 and OT8 in monoculture and co-culture experiments, cell abundance was quantified by SYBR Green-based quantitative PCR (qPCR) targeting the 16S rRNA gene. qPCR was performed on a CFX Connect Real-Time PCR Detection System (Bio-Rad Laboratories, USA) using the PowerUp SYBR Green Master Mix for qPCR (Thermo Fisher Scientific). Total DNA was extracted from cell pellets using the ISOSPIN FECAL kit according to the manufacturer’s instructions. Primer pairs OT_1F (5’-TCTAGCTTGCTAGAGTGGAACT) / OT_1R (5’-CCACTAAAGGGTAGGTTCCTAC) and Mc_2F (5’-GGTGAGTAACACATGGCTG) / Mc_3R (5’-CCTTTCCTCAAGGTCCTCTA) were designed based on the 16S rRNA gene sequences of OT8 and Mc4, respectively. The expected amplicon lengths were 67 bp for OT8 and 65 bp for Mc4. Standard curves were generated from 10-fold serial dilutions of purified PCR products with known concentrations. All reactions, including non-template controls (NTCs), were performed in triplicate. Amplification specificity was confirmed by agarose gel electrophoresis and melting curve analysis, and representative amplicons were verified by Sanger sequencing. All qPCR assays showed no amplification in NTCs or in cultures without added OT8 or Mc4 cells and exhibited amplification efficiencies ≥95% with R² values >0.99.

To evaluate host RNA utilization by OT8 and its impact on host cellular RNA pools, transcript-to-gene copy number ratios were quantified for Mc4 16S rRNA gene and the gene encoding ribosomal protein S12. Total DNA and RNA were extracted from cell pellets using the ISOSPIN FECAL kit and ISOIL for RNA, respectively, following the manufacturer’s protocol. For RNA samples, removal of genomic DNA followed by reverse transcription was performed using the PrimeScript RT reagent kit with gDNA Eraser (TAKARA, Japan) following the manufacturer’s protocol. qPCR was performed for both DNA and RNA (cDNA) samples as described above. For ribosomal protein S12, primer pair S12_3F (5’-GCGTGCGTCTCACTAACCAG) / S12_3R (5’- ACACTGTGCTCCTGTAACCC) was designed based on the Mc4 genome sequence, yielding an expected amplicon length of 56 bp. All qPCR assays showed no amplification in negative controls and exhibited amplification efficiencies ≥98% with R² values >0.99. Transcript-to-gene ratios were calculated to assess relative changes in RNA abundance per genomic copy under monoculture and co-culture conditions. While the ribosomal protein S12 primer set efficiently amplified Mc4 genomic DNA, amplification from Mc4 cDNA was not detected (no signal above negative controls) in all samples analyzed. Consequently, S12 transcript-to-gene ratios were not reported.

### Calculation of the demand of nucleotides and nucleic acids

The total nucleotide demand for OT8 cell replication (nucleotides serving as energy input and biosynthesis precursor) was calculated as follows. For DNA, we calculated based on replication of one chromosome with a length of 972550 bases. For ribosomal RNA and transfer RNA, we assumed a total RNA concentration similar to Escherichia coli (0.1 pg per cell) with rRNA and tRNA comprising 80% and 15% of the total RNA pool. For messenger RNA, we calculated the total concentration based on an estimate of total number of mRNA molecules based on eq. 6 from Lynch^55^ [ (8.837 x 10^3^) x (cell volume)^0.36^], cell volume of 0.006375 μm^3^, low turnover rate on the same order as other bacteria (1 per hour) ^55^, and an average mRNA length of 885 bases. A synthesis cost of 1 ATP per (deoxy)nucleotide was estimated based on the biosynthetic pathways found in the OT8 genome and NDPs as the synthesis precursor. For protein, a per cell abundance was estimated using eq. 5 from Lynch^55^ [ (1.6 x 10^6^) x (cell volume)^0.36^], a low turnover rate on the same order as other bacteria (0.01 per hour) ^55^, and direct usage of amino acids as a precursor for protein synthesis. From this, we further estimated how much host cytosol OT8 would need for cell replication based on a host total RNA concentration similar to E. coli (0.1 pg per µm^3^) with rRNA, tRNA, and mRNA comprising 80%, 15%, and 5% of the total RNA pool. Total RNA, rRNA, tRNA, and mRNA were estimated to be synthesized/consumed at rates consistent with the cell doubling (30 hours) (e.g., RNA content per cell / doubling time). The same were calculated for Mc4 based on an estimated cell volume of 0.588 µm^3^ and doubling time of 18.6 hours.

### Cellular fatty acid analysis

Cellular fatty acid compositions were analyzed as described previously^40^. Briefly, cells from anoxic Mc4 monocultures and Mc4–OT8 co-cultures were harvested during the late exponential growth phase. Fatty acid methyl esters (FAMEs) were prepared and analyzed using the Sherlock Microbial Identification System (version 6.0; Microbial ID, MIDI Inc., USA) in conjunction with the TSBA library database (TSBA6 version 6.20).

## Acknowledgments

We acknowledge the Kanto Natural Gas Development Co., Ltd. for collecting samples at their facilities. We also thank Chiwaka Miyako and Rieko Iwanami for assistance in molecular analyses; Fumie Nozawa, Sanae Yamaoka and Mari Ohara for assistance in cultivation experiments; Eri Hara for assistance in microscopy. This work was supported by JSPS KAKENHI Grant Numbers 17K15183, 23H00387 and 23K18158.

## Author Contributions

T.K. and M.K.N. designed the study. T.K., Y.K., H.T. and M.K.N. wrote the manuscript. T.K. cultured and isolated strains Mc4 and OT8. T.K. and H.T. performed physiological experiments. T.K. and M.K.N. performed bioinformatic analyses. T.K., N.H. and X.Y.M. performed microscopic analyses. All authors reviewed the results and approved the manuscript.

## Competing interests

The authors declare no competing interests.

## Data availability

The genome sequences of strains OT8 and Mc4 has been deposited at DDBJ/ENA/GenBank under the accession JBWBKY010000000 and JBWPUY000000000, respectively. Correspondence and requests for materials should be addressed to T.K.

## Notes

### Competing Interest Statement

The authors have declared no competing interest.

