## Supplementary material for "A representative of a ubiquitous bacterial lineage parasitically feeds on host RNA": Supplmenetary Information

**Supplementary Note 1. Cultivation of ultrasmall parasitic bacterium.**

Earth's subsurface is one of the largest habitats of prokaryotes in both volume and biomass[1] and is known to harbor phylogenetically diverse *Minisyncoccota*/Patescibacteriota[2,3]; thus, in this study we aimed to culture and characterize such organisms from an anoxic saline formation water and sediments derived from deep aquifers in natural-gas deposits[4]. We initially incubated the original environmental samples under anoxic conditions without addition of any organic substrates and, in subcultures, used autoclaved biomass from the mother culture. After 8 years of cultivation, microscopy revealed ultrasmall bacterial cells attached to a filamentous host. Based on amplicon sequencing of the 16S rRNA gene, one population was classified to a bacterial group associated filamentous structure—family *Aggregatilineaceae* in the phylum *Chloroflexota*. We isolated the *Aggregatilineaceae* bacterium, designated as strain Mc4, via dilution-to-extinction combined with deep-agar slant cultivation[5] using yeast extract and the culture supernatant of another bacterium isolated from the same enrichment culture; these components are hereafter referred to as "Mc4 growth substrates" (for details, see Methods). We then inoculated the Mc4 culture with filtered enrichment culture selecting for small cells. Here, we successfully obtained a pure co-culture consisting of the ultrasmall cells, designated as strain OT8, and filamentous Mc4 cells.

**Supplementary Note 2. Growth behavior and host specificity of OT8 toward Mc4**

OT8 could grow in the presence of actively growing Mc4 cells but failed to grow in the presence of non-growing Mc4 or Mc4 cell extracts (Fig. 2c). Supporting these observations, OT8 attaches predominantly to Mc4 cell elongation sites, as indicated by staining of newly synthesized peptidoglycan through amendment of a fluorescent D-amino acid[6] to OT8–Mc4 co-cultures (Supplementary Fig. S2).

We next tested whether active host growth was required only for the initiation of OT8 attachment or for the entire infection process. Cells from a growing Mc4–OT8 co-culture were washed and transferred to fresh medium lacking the substrates required for Mc4 growth. In contrast to the experiment shown in Fig. 2c, in which the OT8 inoculum was prepared by filtration to remove Mc4 cells and Mc4-associated OT8 cells, this washed cell fraction contained OT8 cells already associated with Mc4. Under this condition, OT8 still increased in abundance, presumably because OT8 cells that had already attached to Mc4 before transfer were able to continue growth, although to a lesser extent than in cultures in which Mc4 growth was supported (Supplementary Fig. S3). In contrast, addition of washed Mc4 cells, without Mc4 growth substrates, did not enhance OT8 growth. These results suggest that OT8 initiates attachment specifically to actively growing Mc4 cells, while already attached OT8 cells can continue to grow even when host growth is no longer supported.

Further verifying that OT8 is an obligate parasite dependent on Mc4, physical segregation of planktonic OT8 from Mc4 for more than two weeks severely stunted growth (Supplementary Fig. S4), suggesting that planktonic OT8 likely lose viability if they fail to encounter a host within a certain timeframe. We also examined whether it could parasitize *Aggregatilinea lenta* strain MO-CFX2, the closest cultivated relative of Mc4 (92.5% sequence similarity in the 16S rRNA gene). OT8 neither grew nor attached to MO-CFX2, indicating that OT8 does not parasitize this strain.

**Supplementary Note 3. Host-directed tubular structures at the OT8–Mc4 interface.**

At the interface, we observed tubular structures (14.3±1.4 nm in diameter) connecting OT8 cells with host cells (Fig. 2e and Supplementary Fig. S5a,b). These structures appeared to extend from the attached OT8 cell toward or into the host envelope, suggesting a host-penetrating connection between the parasite and host. Consistent with the presence of these distinct extracellular structures, the OT8 genome encodes homologs of both type IV pili (T4P) [7] and type IV secretion system (T4SS) components (Supplementary Table S1), the latter resembling T4SSs found in *Bacillati* monoderms[8]. The pili-like structure observed in a planktonic OT8 cell (Supplementary Fig. S5c) is likely T4P, which are widely involved in planktonic motility and host attachment[7]. In contrast, the thicker tubular structures observed in host-attached OT8 cells are consistent with T4SS given their larger diameter than typical T4P[9,10] and their apparent penetration into the host envelope, reminiscent of macromolecule-translocating T4SS[11]. Although the transported substrates remain to be identified, the coexistence of host cytoplasmic shrinkage and host-directed tubular structures suggests that OT8 may access host-derived cellular material through direct physical contact.

**Supplementary Note 4. Exclusion of fluorescence derived from transferred free AzG-labeled guanosine or nucleotide pools.**

The AzG-labeling experiment was interpreted as evidence for acquisition of host-derived RNA by OT8 from Mc4. We considered two alternative explanations: first, that AzG-labeled guanosine or guanosine phosphates are transferred from Mc4 to OT8 and directly generate fluorescence within OT8; and second, that transferred guanosine phosphates are used by OT8 as precursors for RNA synthesis, thereby producing AzG-labeled RNA within OT8.

The first possibility is unlikely because free AzG-labeled guanosine and guanosine phosphate derivatives are not expected to produce detectable fluorescence at physiological intracellular concentrations[12]. Thus, the fluorescence signal observed throughout attached OT8 cells cannot be explained simply by intracellular accumulation of soluble AzG9-derived metabolites.

The second possibility is also unlikely on quantitative grounds. Even if AzG-labeled guanosine phosphates were transferred from Mc4 to OT8 and subsequently used as precursors for RNA synthesis within OT8, the soluble free nucleotide pool of a single Mc4 cell is insufficient to support the level of RNA synthesis required for OT8 replication or cell-wide RNA labeling. As evaluated in Supplementary Note 5, free nucleotides account for only a minor fraction of the predicted nucleotide demand of OT8, whereas total RNA and rRNA in Mc4 are sufficient candidate sources. Thus, transfer of soluble AzG-labeled nucleotide precursors alone cannot explain the extensive AzG fluorescence observed throughout attached OT8 cells.

Weak RNA fluorescence was occasionally detected in host-detached planktonic OT8 cells (OT8_P) (Fig. 2d). However, this weak signal was observed only in OT8_P cells located close to Mc4 cells, whereas OT8_P cells spatially separated from Mc4 showed no detectable RNA fluorescence. This spatial association suggests that the weak RNA fluorescence in a subset of OT8_P cells likely reflects optical spillover or local background from adjacent AzG-labeled Mc4 cells, rather than genuine AzG incorporation into OT8-derived RNA.

**Supplementary Note 5. Energetic and biosynthetic sufficiency of host-derived RNA for OT8 replication.**

We evaluated whether host-derived nucleotide pools could meet the energetic and biosynthetic requirements of one OT8 cell division under the reconstructed metabolic constraints described in Methods. The nucleotide demand of OT8 was estimated from chromosome replication, RNA synthesis, and nucleotide-derived energy input, assuming NDPs as biosynthetic precursors and an energetic cost of 1 ATP equivalent per incorporated (deoxy)nucleotide. Host-derived nucleotide availability was then estimated from Mc4 cell volume, RNA content, RNA composition, and doubling time, using total RNA, rRNA, tRNA, mRNA, and free nucleotide pools as candidate sources. The estimated total RNA and rRNA contents of a single Mc4 cell were each sufficient to support one OT8 cell division, whereas tRNA, mRNA, and free nucleotide pools were not. Specifically, tRNA, mRNA, and free nucleotides would satisfy only 49.6%, 16.5%, and 2.4% of the predicted demand, respectively. Rate-based estimates led to the same conclusion: continuous synthesis of total RNA or rRNA by a single Mc4 cell could theoretically sustain replication of at least four OT8 cells, whereas synthesis rates of tRNA, mRNA, or free nucleotides alone were insufficient.

**Supplementary Note 6. Codon-usage complementarity in Minisyncoccota is not**

**explained solely by genomic G+C content.**

*Minisyncoccota*/Patescibacteriota generally have AT-rich genomes, whereas Mc4 and other previously identified hosts have GC-rich genomes. Therefore, the apparent codon-usage complementarity between parasites and hosts could, in principle, arise simply from contrasting genomic G+C contents. To test this possibility, we compared codon usage in ribosomal protein genes between *Minisyncoccota* and non-*Minisyncoccota* bacteria with similar G+C contents. Relative synonymous codon usage (RSCU) profiles were calculated using 33 core ribosomal protein subunits present in ≥97% of genomes, and only genomes encoding at least 30 of these subunits were included to minimize biases caused by incomplete genome recovery. Despite similar genomic G+C contents, *Minisyncoccota* showed distinct ribosomal-protein codon-usage profiles relative to non-*Minisyncoccota* bacteria with OT8-like G+C contents. Several codons were enriched or depleted in *Minisyncoccota* relative to G+C-matched non-Minisyncoccota genomes, based on an effect-size criterion and statistical support (Supplementary Fig. S11). Within-class correspondence analysis of ribosomal-protein RSCU profiles further separated *Minisyncoccota* from non-*Minisyncoccota* genomes in multivariate space, with WCA axes 1–3 explaining 61.6% of the variance and significant separation by PERMANOVA ($p$ = 0.001; Supplementary Fig. S12). These results indicate that *Minisyncoccota* codon usage is not merely a passive consequence of low genomic G+C content.

We next asked whether this *Minisyncoccota*-specific codon usage corresponds to reduced predicted expressivity of host-derived mRNAs in *Minisyncoccota* cells. Using ribosomal protein genes from each parasite genome as the translational reference set, we calculated MELP[13]-based expression-potential scores for host genes as target sequences. Among bacteria with OT8-like G+C contents, OT8 and other *Minisyncoccota* showed lower codon-usage compatibility with genes from Mc4 and other previously identified hosts than did non-*Minisyncoccota* taxa (Fig. 4b and Supplementary Fig. S13). Conversely, when genes from bacteria with Mc4-like G+C contents were evaluated under three *Minisyncoccota* translational backgrounds, representatives of *Minisyncoccota*, *i.e.*, OT8, *Minisyncoccia*, and *Ca.* Dojkabacteria genomes all showed broadly low expression potential for these GC-rich genes (Supplementary Fig. S14).

Together, these analyses indicate that codon-usage complementarity in *Minisyncoccota* is both lineage-specific and broad: it distinguishes *Minisyncoccota* from G+C-matched non-*Minisyncoccota* bacteria and extends beyond the OT8–Mc4 pair to genes from previously identified hosts and phylogenetically diverse GC-rich bacteria. This pattern supports the interpretation that complementary codon utilization is not simply a consequence of contrasting genomic G+C contents, but may reflect a broader evolutionary relationship between *Minisyncoccota* codon usage and exposure to host-derived RNA.

### Supplementary Note 7. Host-attached morphology and polarized cell division of OT8.

Cryo-electron microscopy (CryoEM) revealed that OT8 cells adopted distinct morphologies depending on their association with host cells (Supplementary Fig. S17). Planktonic OT8 cells were tapered and bullet-like, with a single pili-like wavy filament on the flatter side of the cell opposite the tapered pole (5.9 ± 0.3 nm in diameter; Supplementary Fig. S5b). This filament-bearing flatter side corresponded to the side by which OT8 cells landed on and attached to host cells. Upon attachment, OT8 cells became rounder and substantially larger, showing a 4.6-fold increase in cytoplasmic volume.

These observations indicate that OT8 undergoes marked morphological remodeling upon host attachment.

Accurate measurements of OT8 cell volumes using CryoEM revealed that the cytoplasmic volume of parasitizing mother cells ranged between 0.023 to 0.037 $\mu m^3$. The interquartile range of the volumes was 0.0066 $\mu m^3$, remarkably close to the average cytoplasmic volume of OT8 daughter cells (0.0064 ± 0.00061 $\mu m^3$). We suggest that OT8 cell first expands, and that this additional cell volume is then used to form a daughter cell through polar division. The resulting daughter cells faced the opposite direction, with their filament-bearing landing side oriented away from the host-attached mother cell (Supplementary Fig. S17). Thus, host-associated cell enlargement is spatially coupled to polarized, directional cell division.

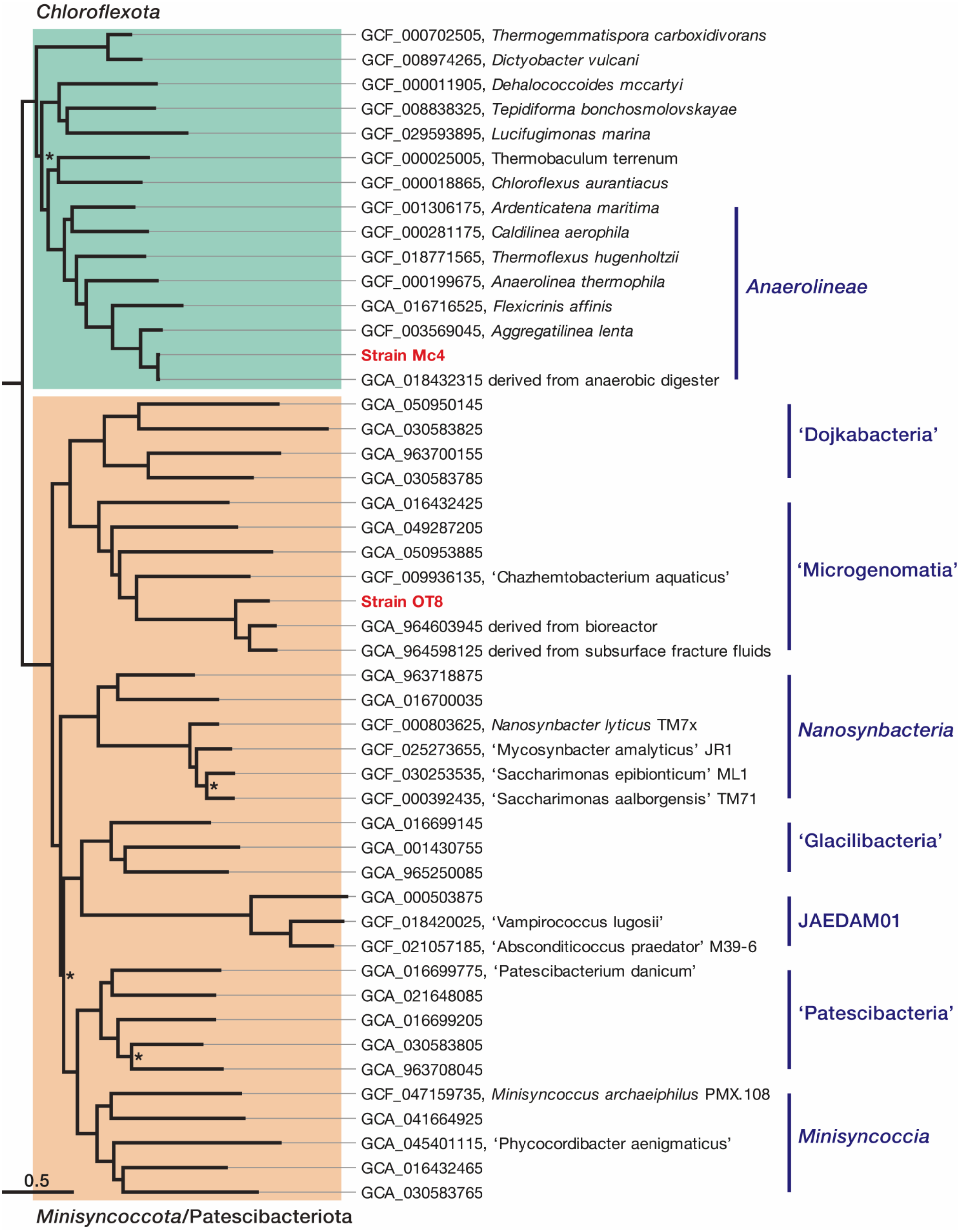


**Supplementary Fig. S1 | Phylogenetic placement of OT8 and Mc4 within the phyla *Minisyncoccota*/Patescibacteriota and *Chloroflexota*.**
Maximum-likelihood phylogeny of representative *Minisyncoccota*/Patescibacteriota and *Chloroflexota* genomes inferred from a concatenated alignment of conserved marker proteins. The tree was rooted using *Actinomycetota* sequences (not shown). Taxonomic labels on the right indicate major classes. Asterisks at internal nodes indicate branches with ultrafast bootstrap approximation support <95% and SH-like approximate likelihood ratio test support <80%, respectively.

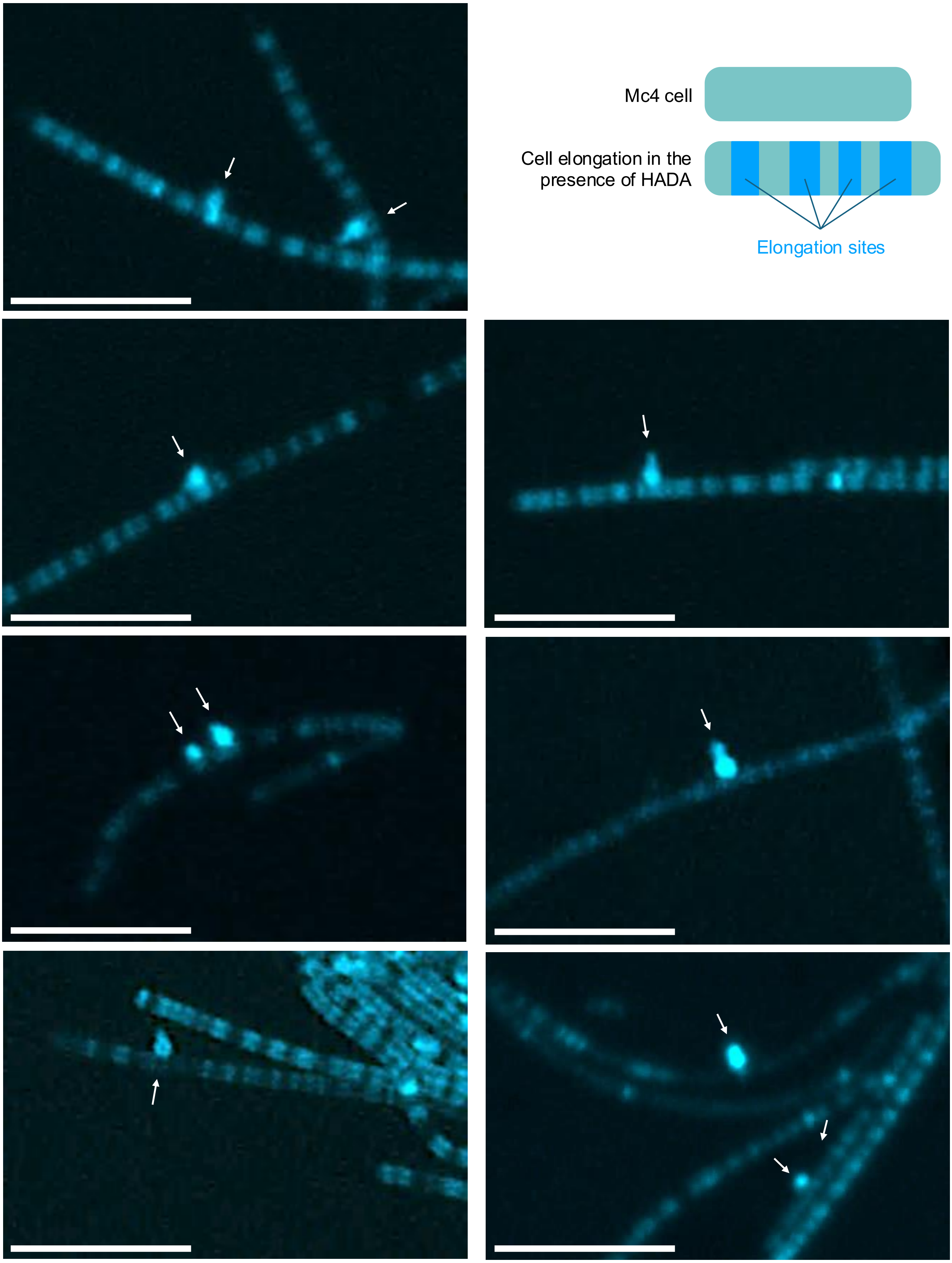


**Supplementary Fig. S2 | Fluorescence microscopy image and schematic illustration of OT8 cells attached to elongation sites of Mc4 cells.**
Strain OT8 and the fluorescent D-amino acid HADA were added to a culture containing growing Mc4 and OT8 cells and incubated for 2.5 days. HADA incorporation marks sites of active cell-wall synthesis in elongating Mc4 cells, which are shown as bright cyan regions in the fluorescence microscopy image and schematically indicated as blue segments in the illustration. White arrows indicate OT8 cells attached near HADA-positive elongation sites on the Mc4 cell surface. Due to the short incubation period, only one or two OT8 cells were observed attached to individual Mc4 cells. Scale bar, 5.0 µm.

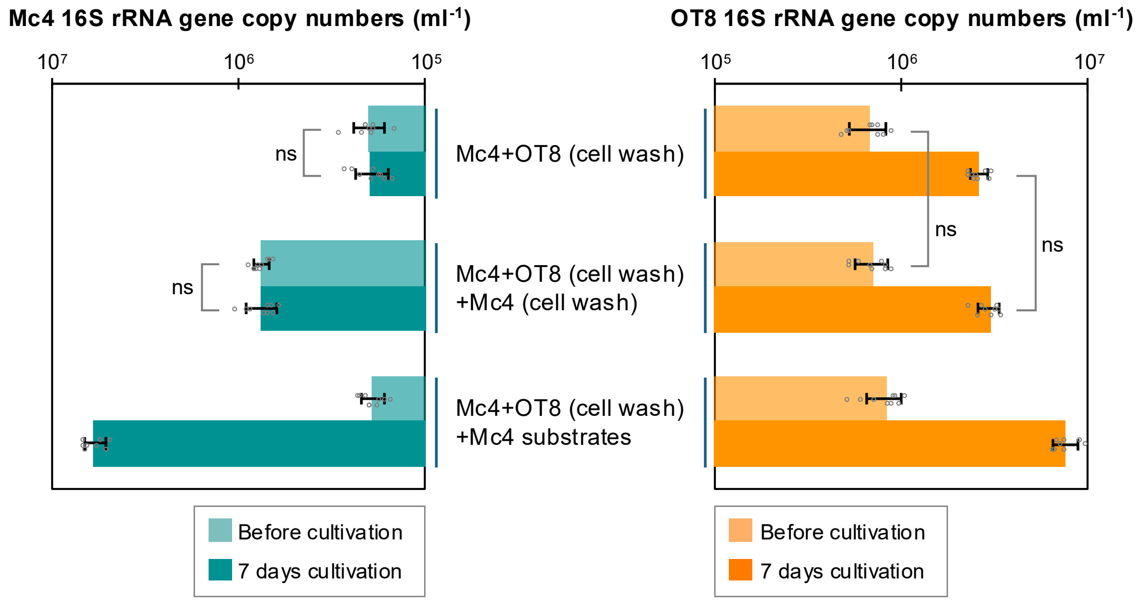


**Supplementary Fig. S3 | Parasitic growth of OT8 on actively growing host Mc4.**
16S rRNA gene copy numbers of Mc4 (left) and OT8 (right), measured at the start and after 7 days of cultivation under three conditions. Washed cells from a Mc4–OT8 co-culture were incubated (i) in fresh medium without Mc4 growth substrates (top), (ii) together with washed cells from an Mc4 axenic culture in fresh medium without Mc4 growth substrates (middle), or (iii) in fresh medium containing Mc4 growth substrates (bottom). Data represent the mean of triplicate experiments, with relative s.d. shown as error bars. Statistical significance was evaluated using a t-test. Abbreviation: ns, not significant.

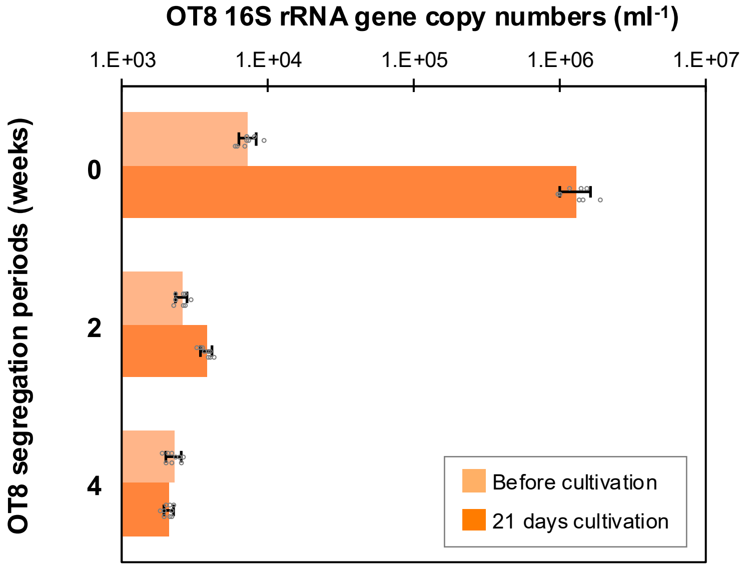


**Supplementary Fig. S4 | Dependence of OT8 on Mc4.**
16S rRNA gene copy numbers of OT8 in co-cultures inoculated with Mc4 cells and host-detached OT8 cells that were collected via filtration and incubated for 0, 2, or 4 weeks in the absence of its host.

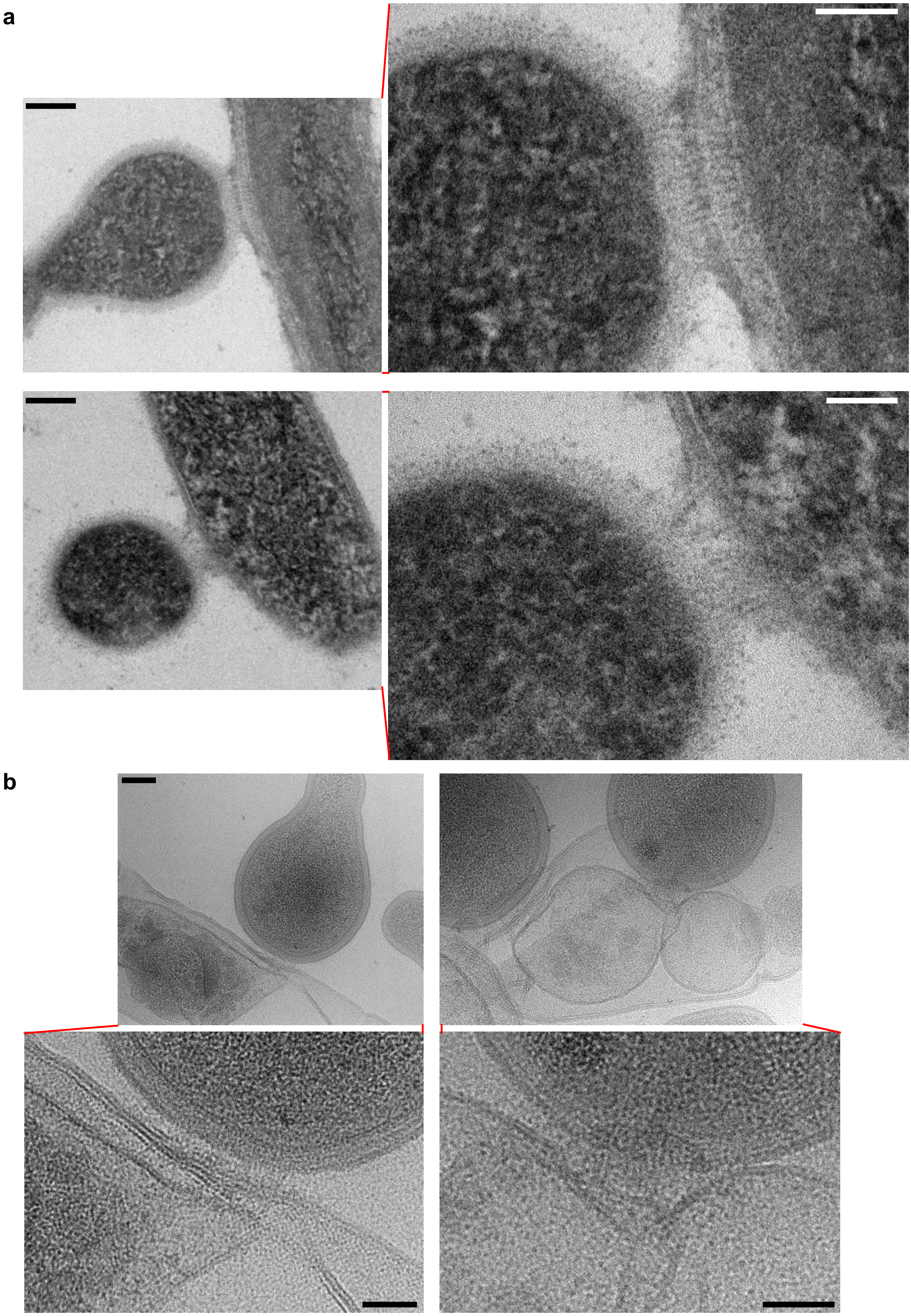


**Supplementary Fig. S5 | Pili-like structures in OT8 cells.**
**a**,**b**, Transmission electron micrographs of ultrathin sections (**a**) and cryo-electron micrographs (**b**) showing tubular connections between OT8 and Mc4 cells. Scale bars: 100 nm (**a**, **b**); 50 nm (enlarged views in **a**, **b**). (continued on next page)

c

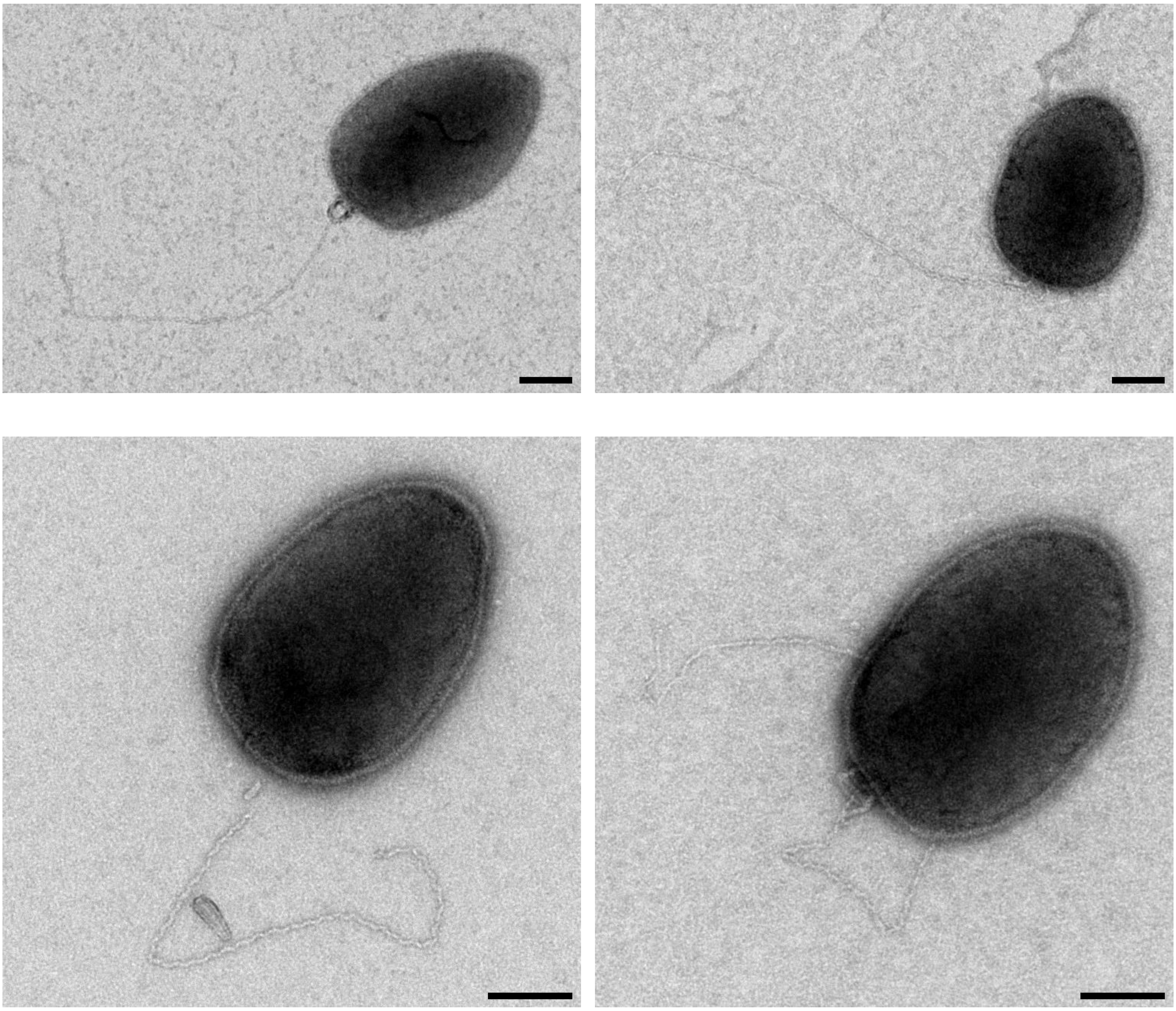

**Supplementary Fig. S5 (continued) | Pili-like structures in OT8 cells.**
**c**, Negative-stain TEM image showing a pili-like structure on planktonic OT8. Scale bars: 100 nm (**c**).

a

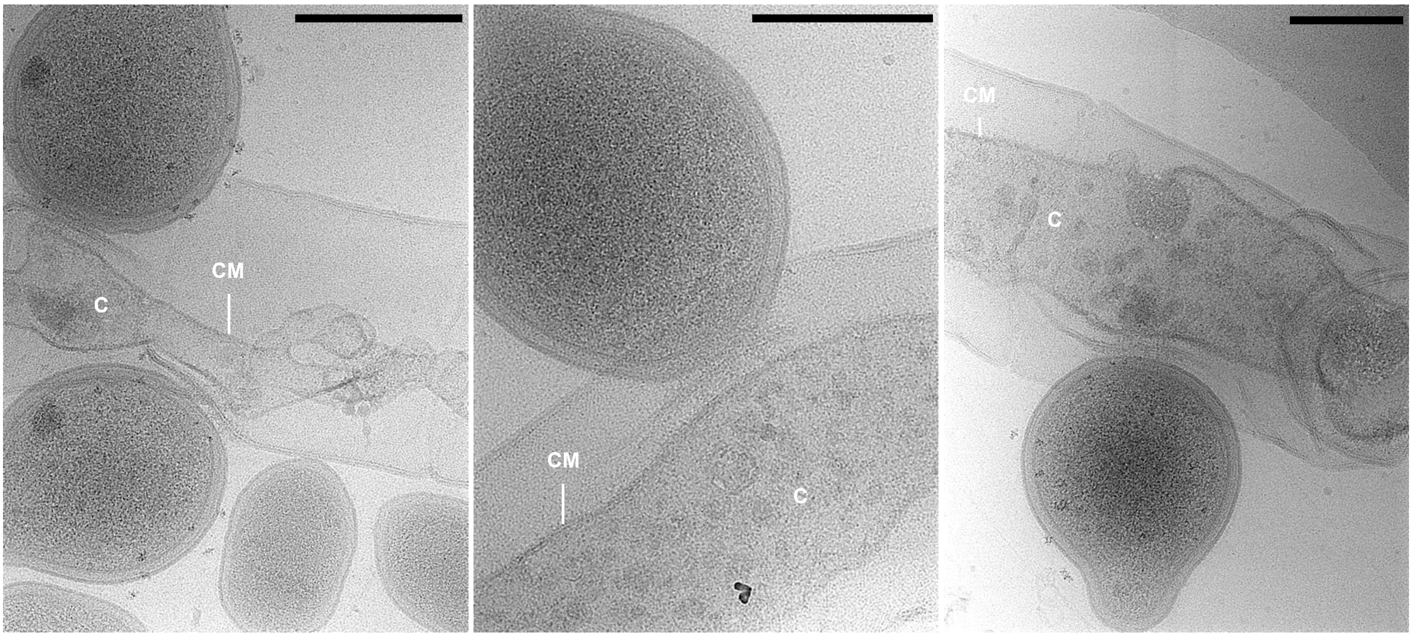


b

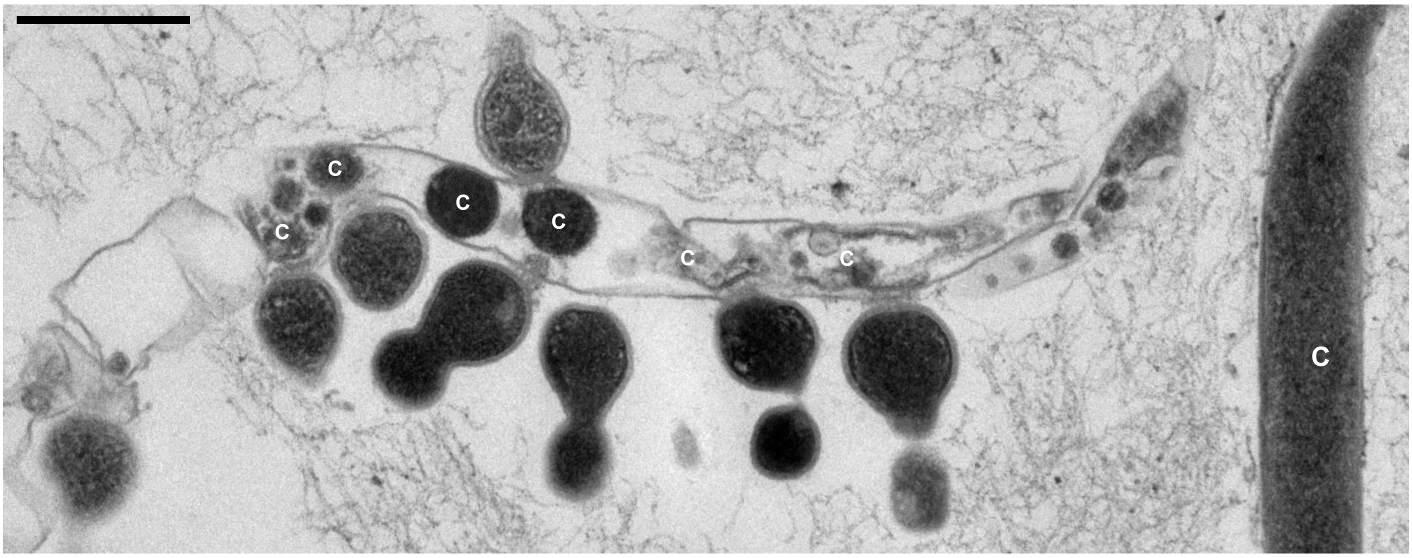


**Supplementary Fig. S6 | Cytoplasmic shrinkage in OT8-infected Mc4 cell cytoplasm.**
**a**,**b**, Cryo-electron micrographs (**a**) and transmission electron micrograph of ultrathin sections (**b**) showing Mc4 cells infected by OT8. Abbreviations: C, cytoplasm; CM, cytoplasmic membrane. Scale bars: 200 nm (**a**); 500 nm (**b**).

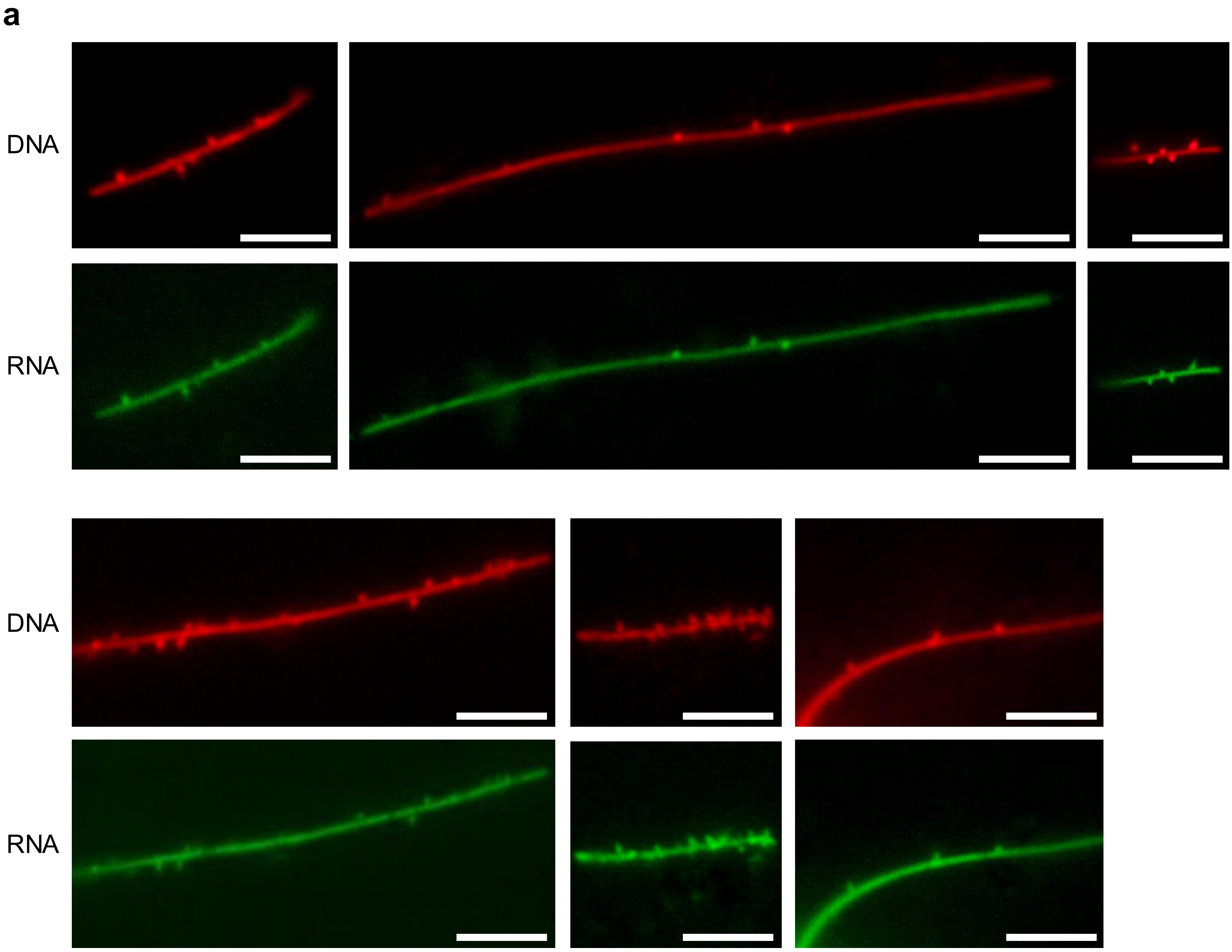


**Supplementary Fig. S7 | Fluorescent labeling of DNA and RNA in OT8 and Mc4 cells.**
**a**, Representative fluorescent micrographs of cells from Mc4–OT8 co-cultures. DNA and RNA were stained with SYTO59 and AzG, respectively. Scale bars: 5 µm. (continued on next page)

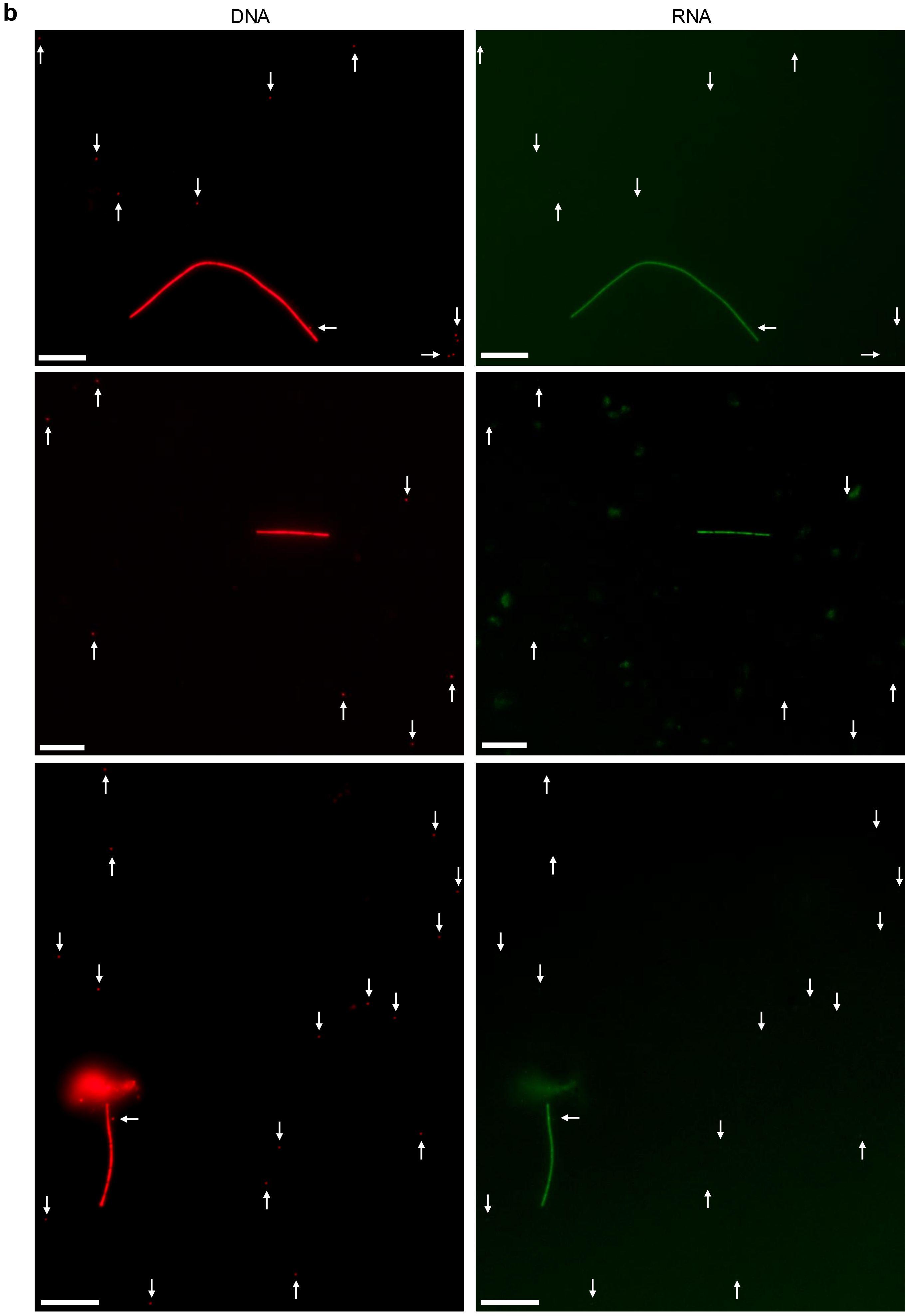


**Supplementary Fig. S7 (continued) | Fluorescent labeling of DNA and RNA in OT8 and Mc4 cells.** **b**, Representative fluorescent micrographs of cells from a mixture of Mc4 monocultures and Mc4-free OT8 cells. DNA and RNA were stained with SYTO59 and AzG, respectively. Arrows indicate Mc4-free planktonic OT8 cells. Scale bars: 10 μm.

**a**

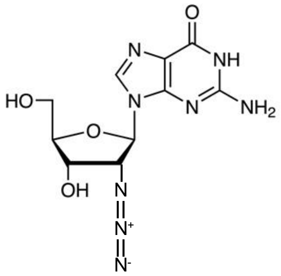


**b**

Guanosine ⟷ (gsk) GMP ⟷ (gmk) GDP ⟷ (ndk) GTP

**c**

| Gene | Protein | OT8 genome | Mc4 genome |
|---|---|---|---|
| gsk | Guanosine-inosine kinase | - | Mc4_04603 |
| gmk | Guanylate kinase | - | Mc4_04106 |
| ndk | Nucleoside diphosphate kinase | OT_00387 | Mc4_04304 |

**Supplementary Fig. S8 | Metabolic capacity underlying the incorporation of AzG into RNA.**
**a**, Chemical structure of 2′-deoxy-2′-azidoguanosine (AzG), a guanosine analog, which was used for microscopic RNA labeling experiments. **b**, Biosynthetic pathways for guanosine triphosphate (GTP) from guanosine. **c**, Presence or absence of genes encoding enzymes for these pathways in the genomes of OT8 and Mc4.

| *Minisyncoccota* ( ) | Host ( ) | G+C% | RNAc. | EP |
|---|---|---|---|---|
| Strain OT8 | Strain Mc4 | 40.8 | + | -2.3 |
| *Ca*. Vampirococcus lugosii | *Halorhodospira halophila* | 21.4 | + | -3.0 |
| *Ca*. Absconditicoccus praedator M39-6 | *Halochromatium roseum* | 27.9 | + | -2.5 |
| *Ca*. Southlakia epibionticum ML1 | *Actinomyces israelii* | 49.1 | + | -1.6 |
| *Nanosynbacter lyticus* TM7x | *Actinomyces odontolyticus* | 44.5 | - | -0.9 |
| *Ca*. Mycosynbacter amalyticus JR1 | *Gordonia amarae* | 50.4 | - | -0.9 |
| Strain TM7i | *Leucobacter aridicollis* | 47.7 | - | -0.6 |
| *Minisyncoccus archaeophilus* PMX.108 | *Methanospirillum hungatei* | 36.4 | - | -0.6 |

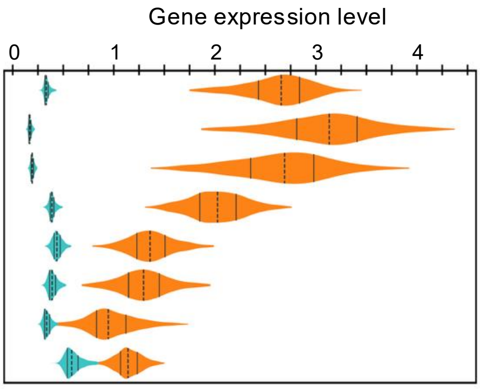


**Supplementary Fig. S9 | Predicted expression potential of host-derived mRNAs in *Minisyncoccota*/Patescibacteriota cells.**

MELP-based expression potential of host-derived mRNAs in *Minisyncoccota*/Patescibacteriota cells. Violin plots show the distributions of predicted expression levels for endogenous *Minisyncoccota* mRNAs (orange) and host-derived mRNAs (green); dotted vertical lines indicate the median and solid vertical lines indicate the first and third quartiles. The host mRNA expression potential (EP) indicates the difference between the median expression potential of host-derived mRNAs and that of endogenous *Minisyncoccota* mRNAs. "G+C%" indicates the genomic G+C content of each *Minisyncoccota* member, and "RNAc." indicates the presence (+) or absence (−) of predicted RNA-catabolic capacity.

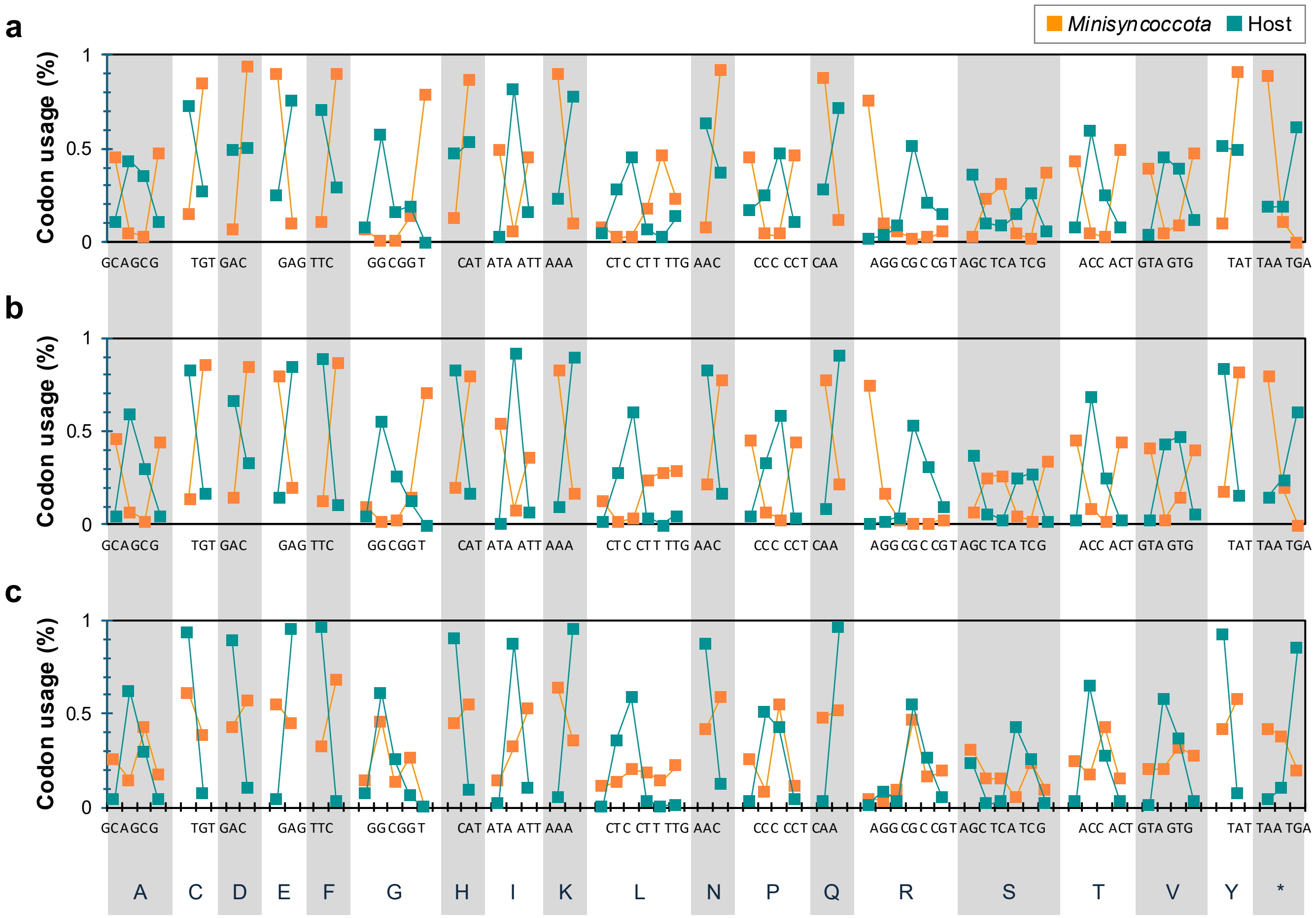


**Supplementary Fig. S10 | Codon usage complementation between putative RNA-degrading *Minisyncoccota*/Patescibacteriota and their hosts.**

**a-c**, Codon usage, shown as the average proportion of total codons per amino acid, shown for the following parasite-host pairs: *Ca.* Vampirococcus lugosii and *Halorhodospira halophila* (**a**), *Ca.* Absconditicoccus praedator M39-6 and *Halochromatium roseum* (**b**), and *Ca.* Southlakia epibionticum ML1 and *Actinomyces israelii* (**c**). As the host genome sequence of *Ca.* V. lugosii is unavailable, the closely related species *Halorhodospira halophila*, identified based on 16S rRNA gene similarity, was used instead (**a**).

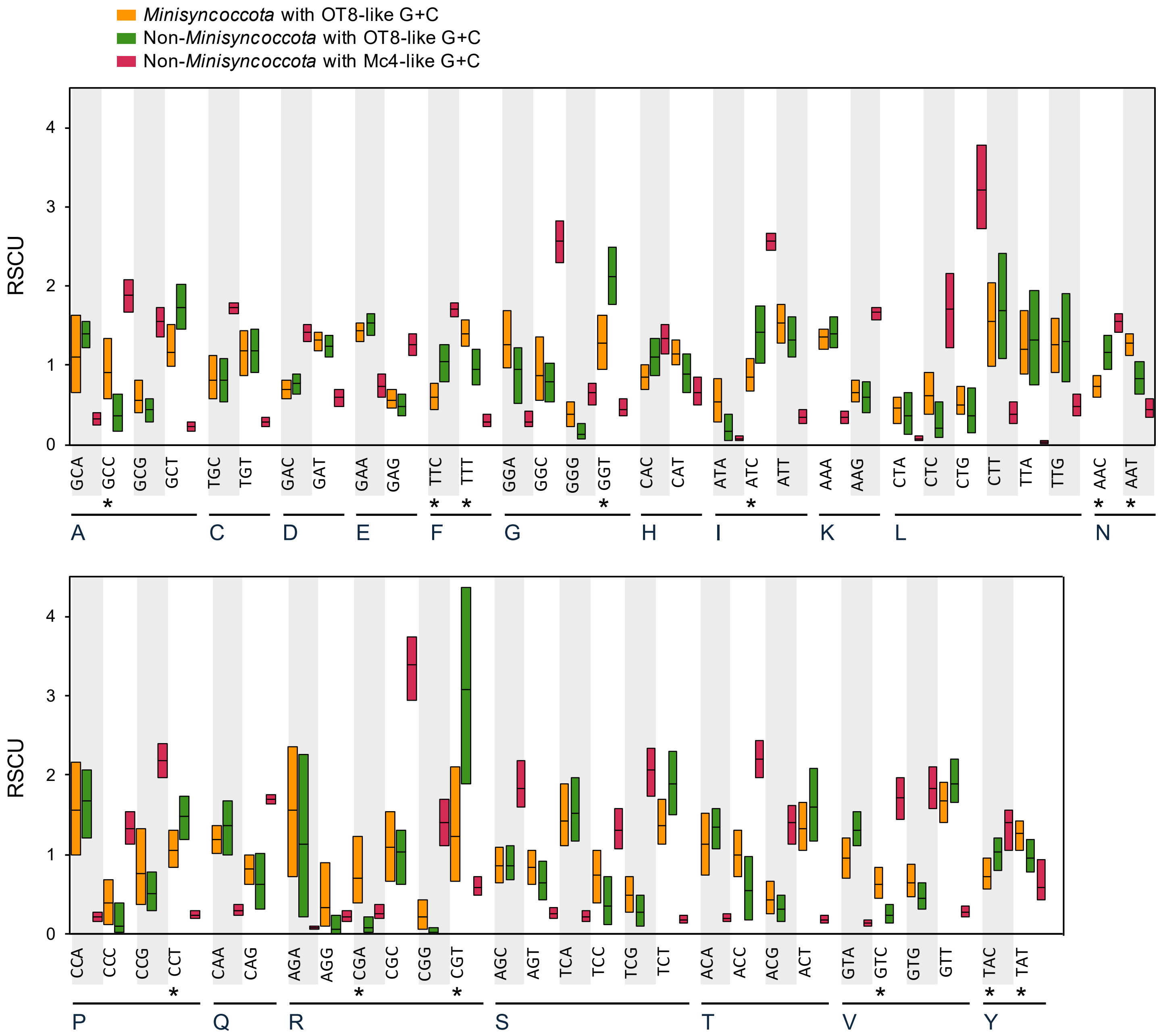


**Supplementary Fig. S11 | Codon usage of ribosomal protein genes in *Minisyncoccota*/Patescibacteriota and other bacteria with OT8-like and Mc4-like G+C contents.** Box plots compare codon-wise relative synonymous codon usage (RSCU) in ribosomal protein–coding genes among *Minisyncoccota* with OT8-like G+C contents (39.3–42.3%), non-*Minisyncoccota* genomes with OT8-like G+C contents, and non-*Minisyncoccota* genomes with Mc4-like G+C contents (63.9–66.9%). Boxes indicate the median and interquartile range. To minimize biases arising from incomplete genome recovery, only genomes encoding at least 30 of a set of 33 core ribosomal protein subunits were included in the analyses. Asterisks mark codons that were selected as *Minisyncoccota*-enriched/depleted relative to non-*Minisyncoccota* genomes with OT8-like G+C contents using an effect-size–based criterion (S_score ≥ 0.8 and |ΔRSCU| ≥ 0.3), and supported by a Kruskal–Wallis test ($p \leq 0.001$).

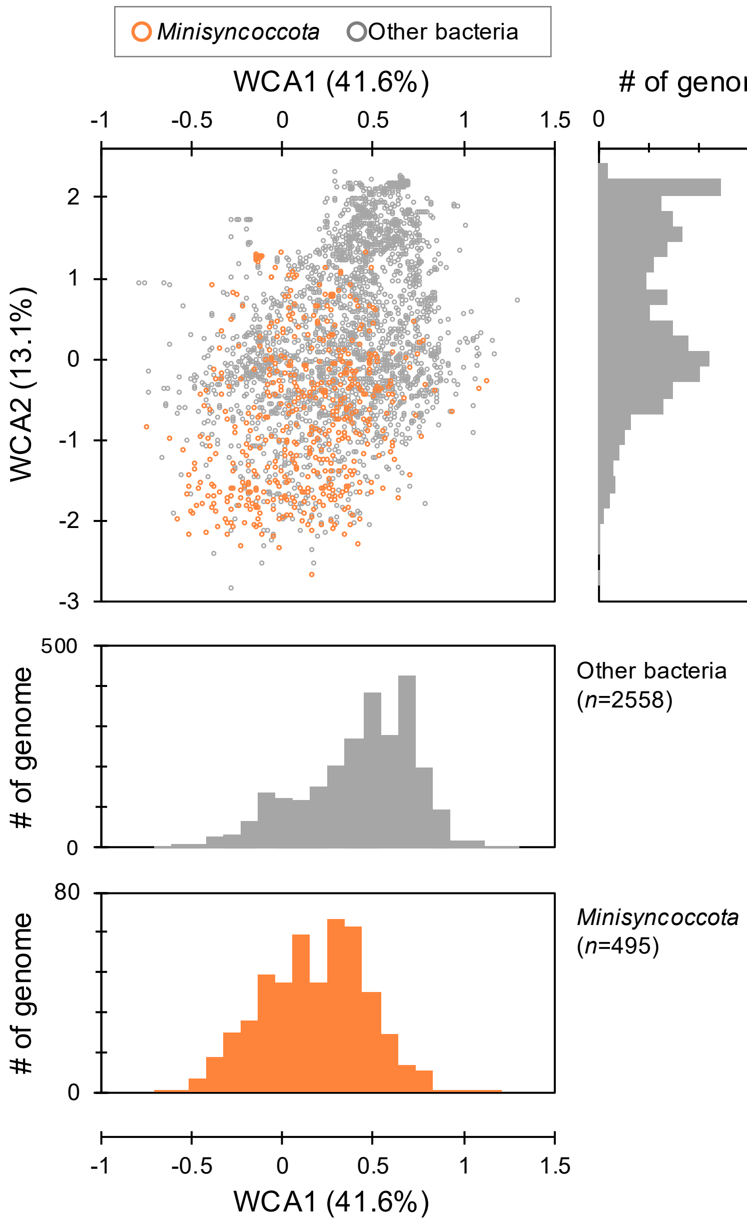


**Supplementary Fig. S12 | Within-class correspondence analysis of ribosomal-protein codon profiles distinguishes *Minisyncoccota*/Patescibacteriota from other bacterial genomes within the OT8-like G+C range.**

Within-class correspondence analysis (WCA) ordination of ribosomal-protein relative synonymous codon usage (RSCU) profiles for individual genomes within the OT8-like G+C range (39.26–42.26%). To minimize biases arising from incomplete genome recovery, only genomes encoding at least 30 of a set of 33 core ribosomal protein subunits were included in the analyses. The upper left panel shows the first two axes, WCA1 and WCA2, each with percentages of inertia of 41.6% and 13.1%. Histograms indicate the distribution of genomes along WCA2 (upper right of the ordination) and WCA1 (lower panels), separately for non-*Minisyncoccota* (gray) and *Minisyncoccota* (orange) genomes. *Minisyncoccota* and non-*Minisyncoccota* genomes were significantly separated in multivariate WCA space (axes 1–3, which together explained 61.6% of the variance), with both PERMANOVA and PERMDISP yielding $p = 0.001$ (9,999 permutations).

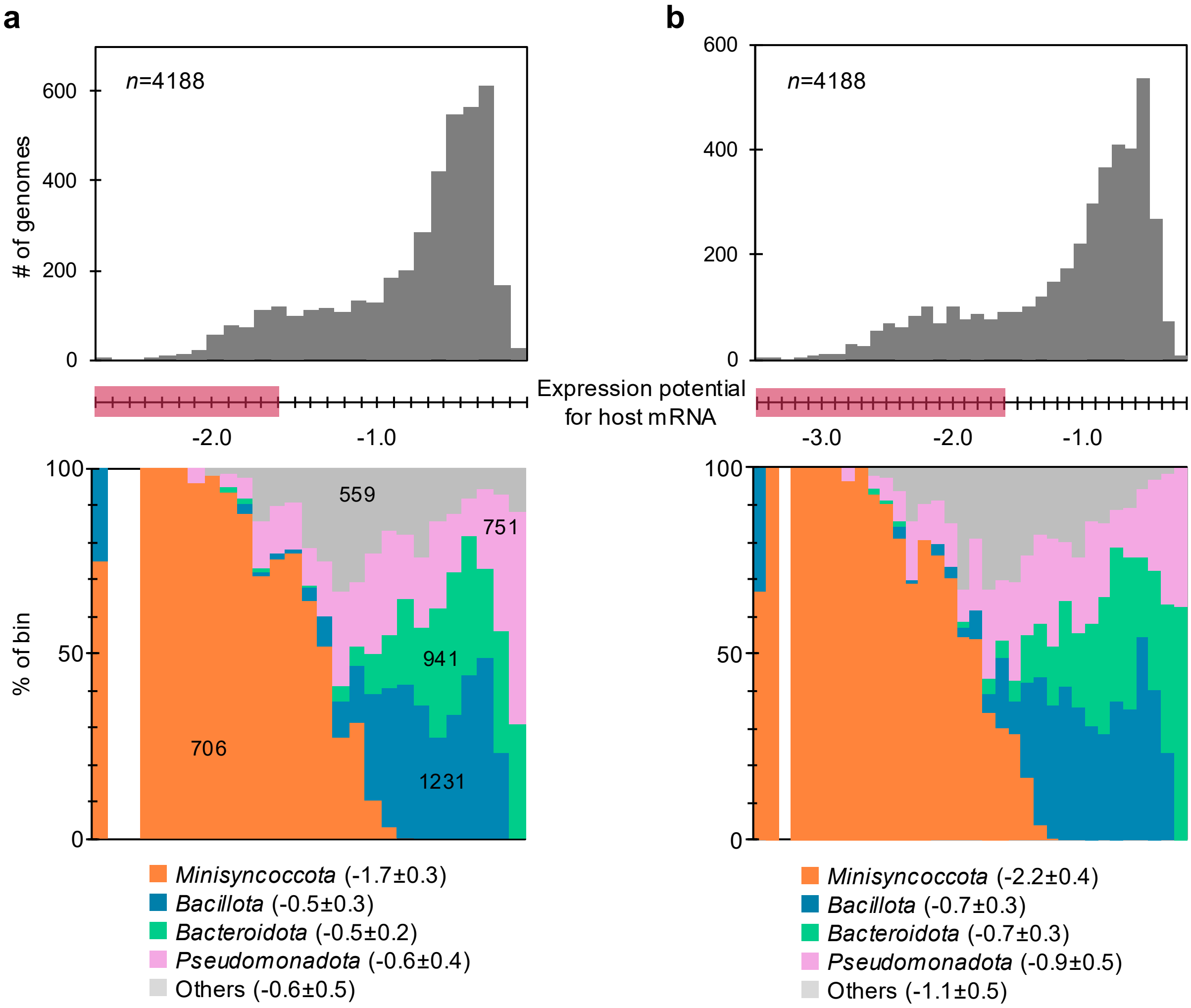


**Supplementary Fig. S13 | Expression potential of identified *Minisyncoccota*/Patescibacteriota hosts against OT8 and other bacterial genomes with OT8-like G+C contents (39.8–43.8%).**
**a**,**b**, Distributions of expression potential for mRNAs of *Minisyncoccota* hosts (*Halochromatium roseum* M39-5 (**a**) or *Actinomyces israelii* F0345 (**b**)) by OT8 and bacteria with genomic G+C contents similar to OT8 (39.8–43.8%). The red range in x-axis corresponds to the range of expression potential index observed in Supplementary Fig. S9 for host-CPR pairs for which RNA catabolism was predicted. Values in parentheses indicate median ± SD expression potentials.

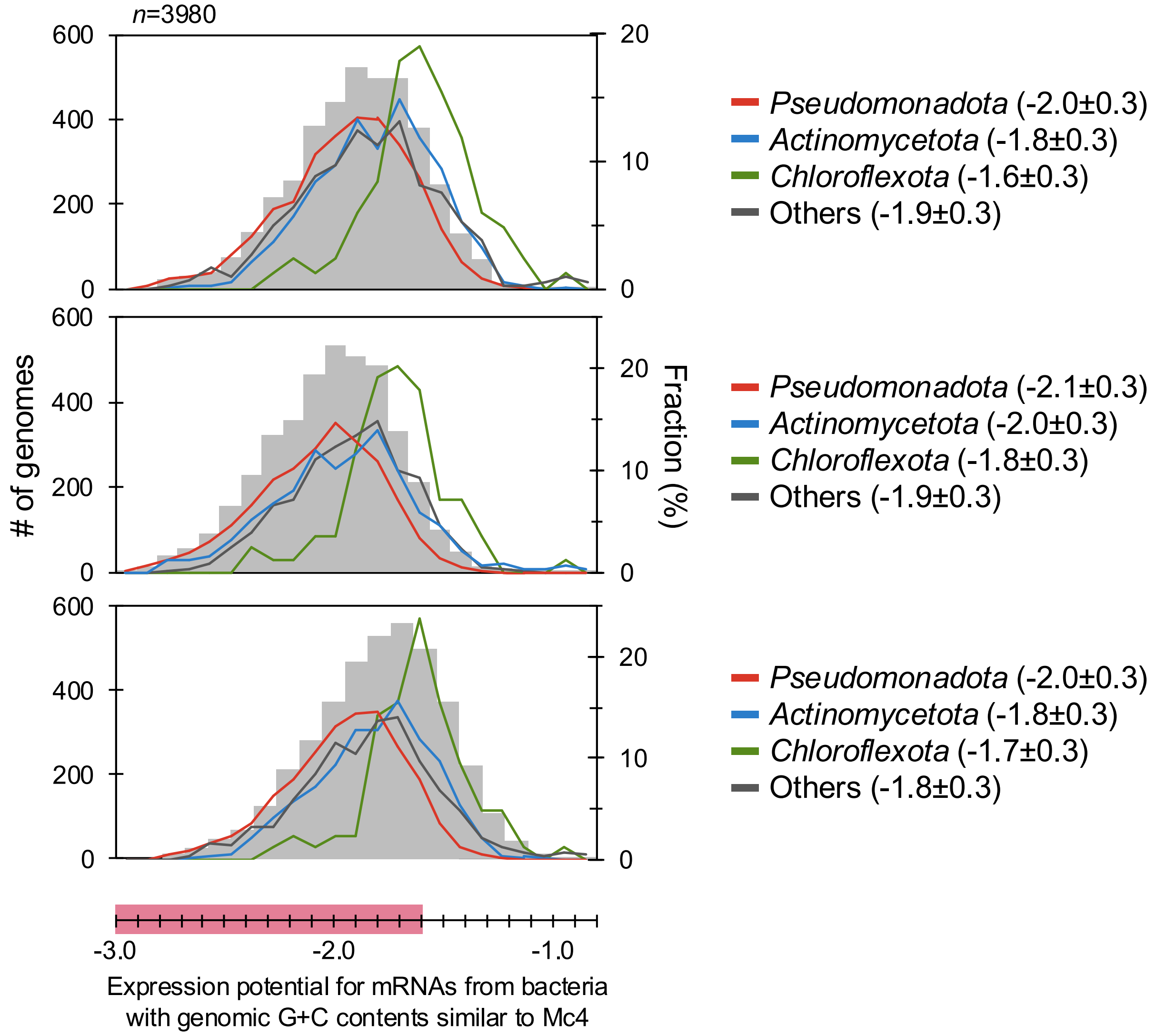


**Supplementary Fig. S14 | Broad codon-usage complementarity between *Minisyncoccota*/Patescibacteriota and GC-rich bacterial lineages.**

Distributions of expression potential for mRNAs from bacteria with with genomic G+C contents similar to Mc4 (63.4–67.4%) against OT8 (upper), *Minisyncoccia* genome (GCA_021734905; 41.1% G+C, middle), and *Ca.* Dojkabacteria genome (GCA_016932785; 40.5% G+C, lower). The red range in x-axis corresponds to the range of expression potential index observed in Supplementary Fig. S9 for host-CPR pairs for which RNA catabolism was predicted. Values in parentheses indicate median ± SD expression potentials.

**a**

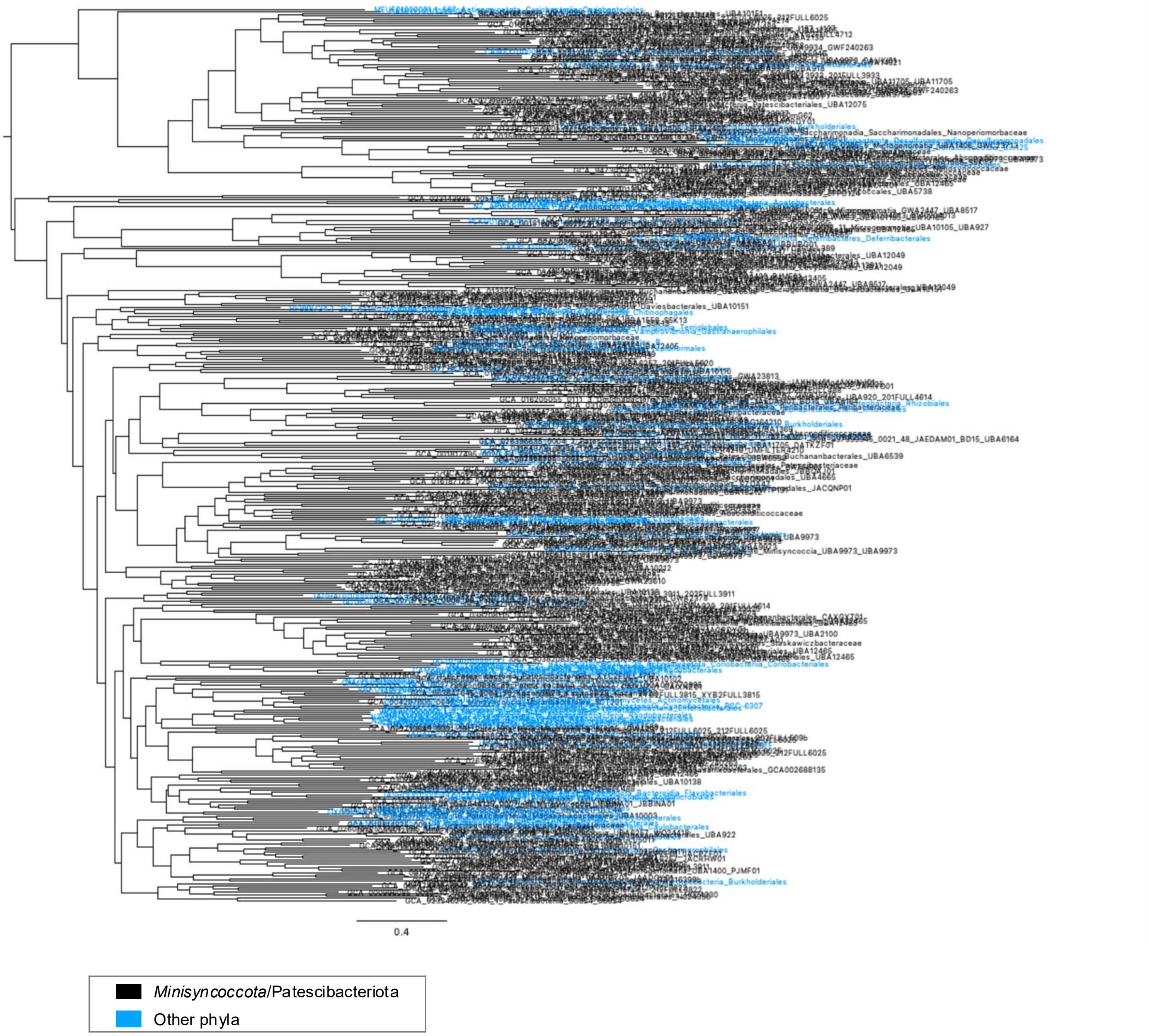


**Supplementary Fig. S15 | Phylogenetic trees of enzymes involved in nucleotide synthesis showing horizontal transfer of these genes.**

**a**, Maximum-likelihood tree estimated from an alignment of purine biosynthesis protein adenylosuccinate synthase. (continued on next page)

**b**

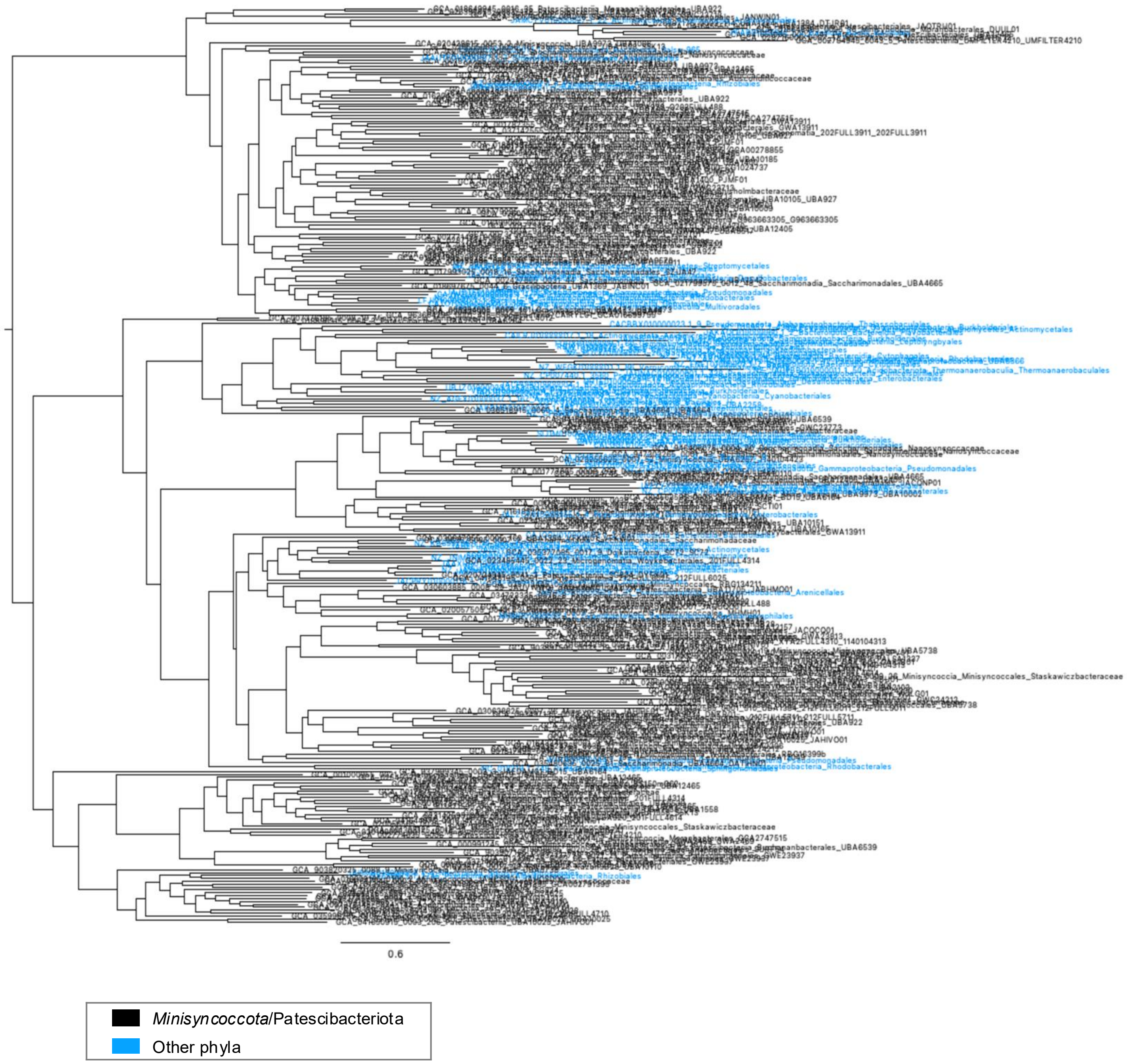


**Supplementary Fig. S15 | Phylogenetic trees of enzymes involved in nucleotide synthesis showing horizontal transfer of these genes.**

**b**, Maximum-likelihood tree estimated from an alignment of purine biosynthesis protein adenylosuccinate lyase. (continued on next page)

**c**

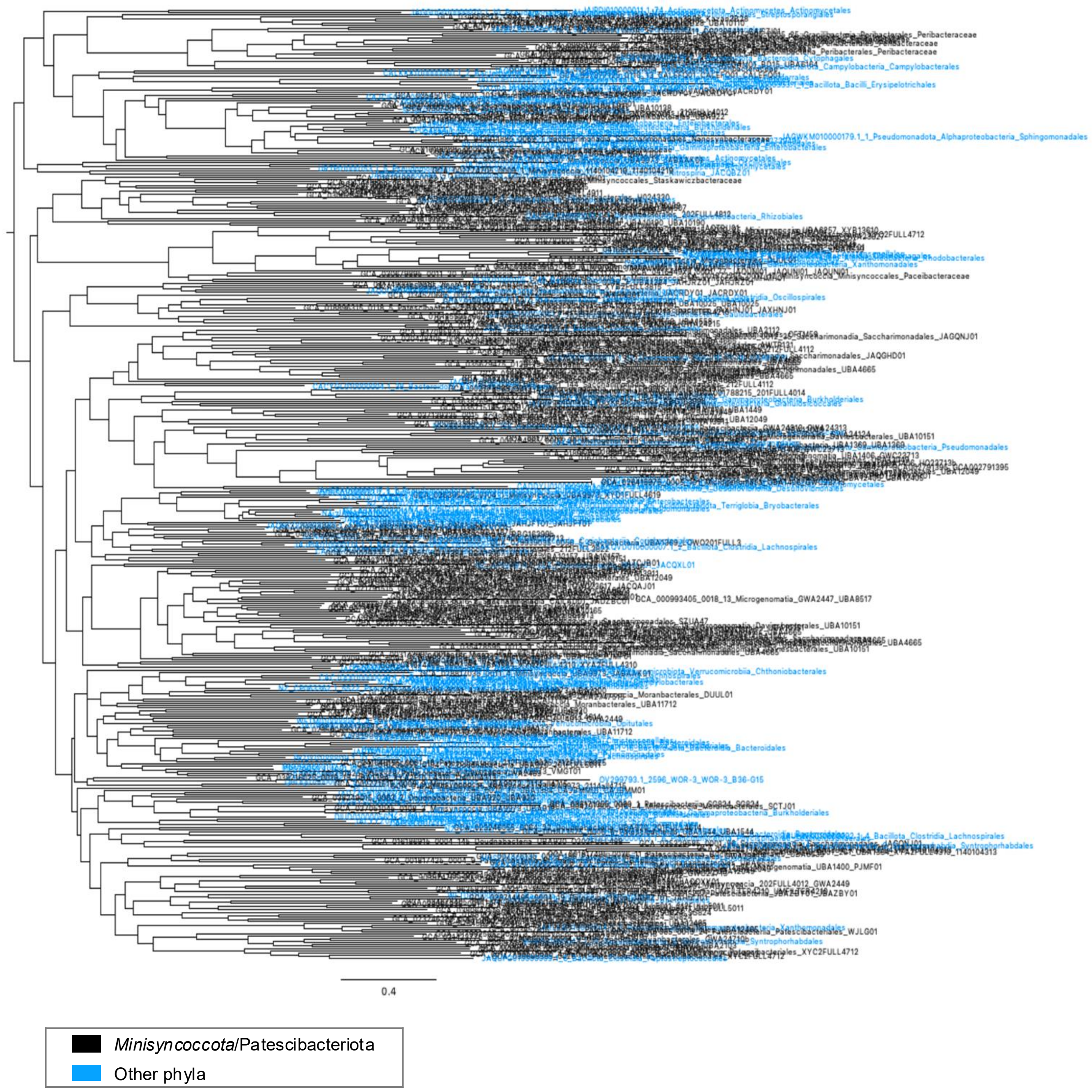


**Supplementary Fig. S15 | Phylogenetic trees of enzymes involved in nucleotide synthesis showing horizontal transfer of these genes.**

**c**, Maximum-likelihood tree estimated from an alignment of purine biosynthesis protein amidophosphoribosyltransferase. (continued on next page)

**d**

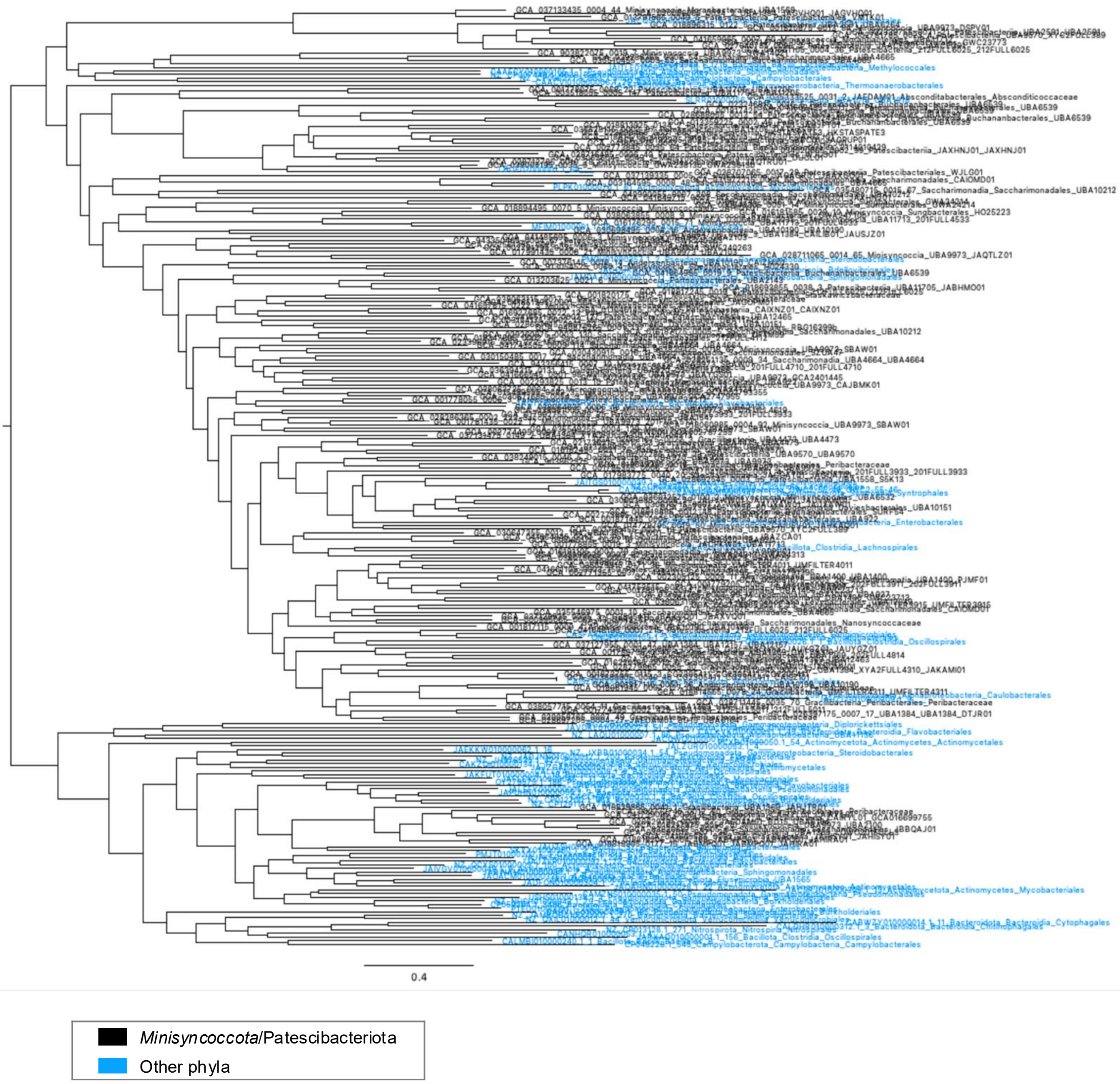


**Supplementary Fig. S15 | Phylogenetic trees of enzymes involved in nucleotide synthesis showing horizontal transfer of these genes.**

**d**, Maximum-likelihood tree estimated from an alignment of pyrimidine biosynthesis protein aspartate carbamoyltransferase. (continued on next page)

**e**

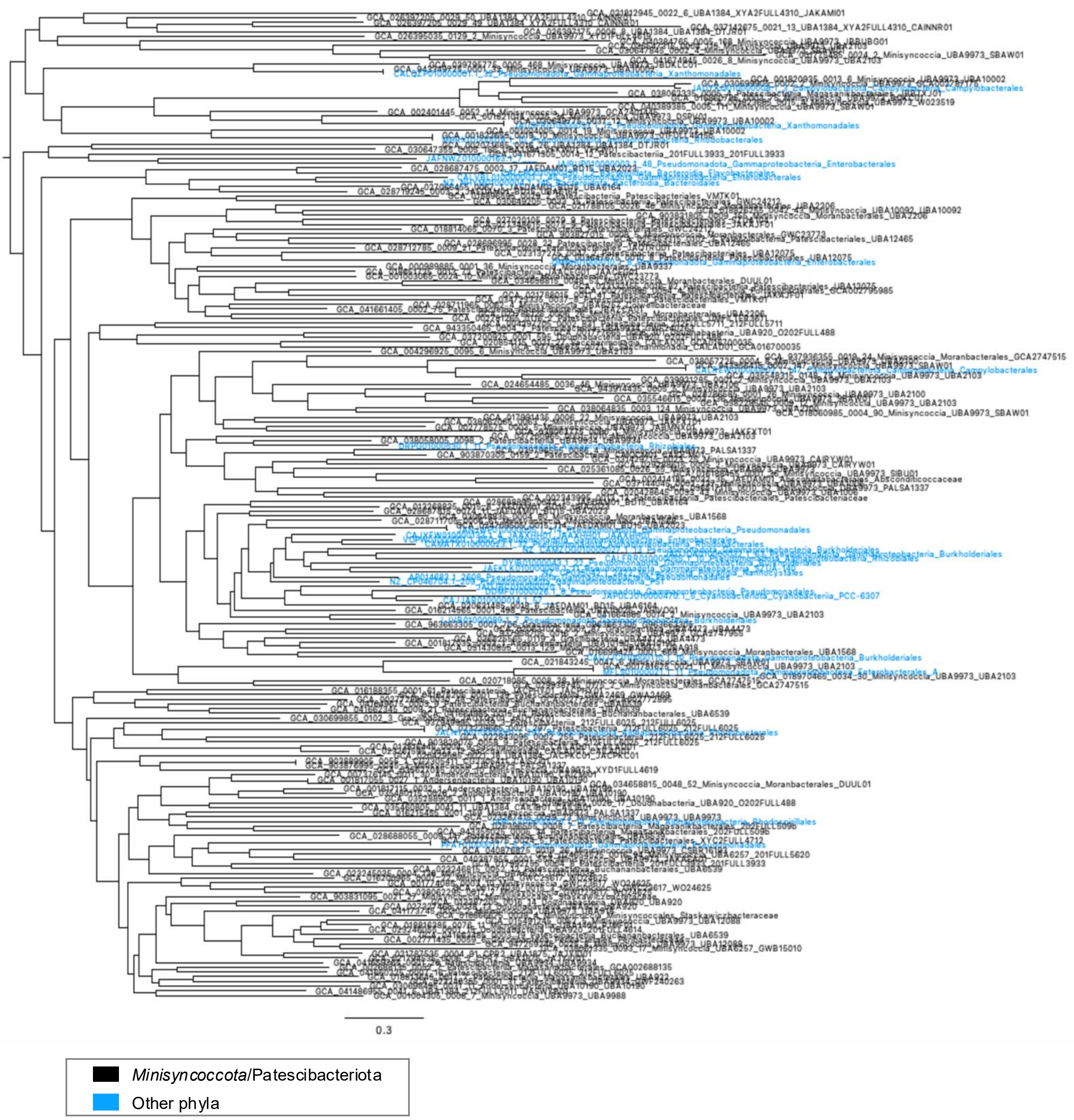


**Supplementary Fig. S15 | Phylogenetic trees of enzymes involved in nucleotide synthesis showing horizontal transfer of these genes.**

**e**, Maximum-likelihood tree estimated from an alignment of pyrimidine biosynthesis protein aspartate dihydroorotase (COG0418). (continued on next page)

f

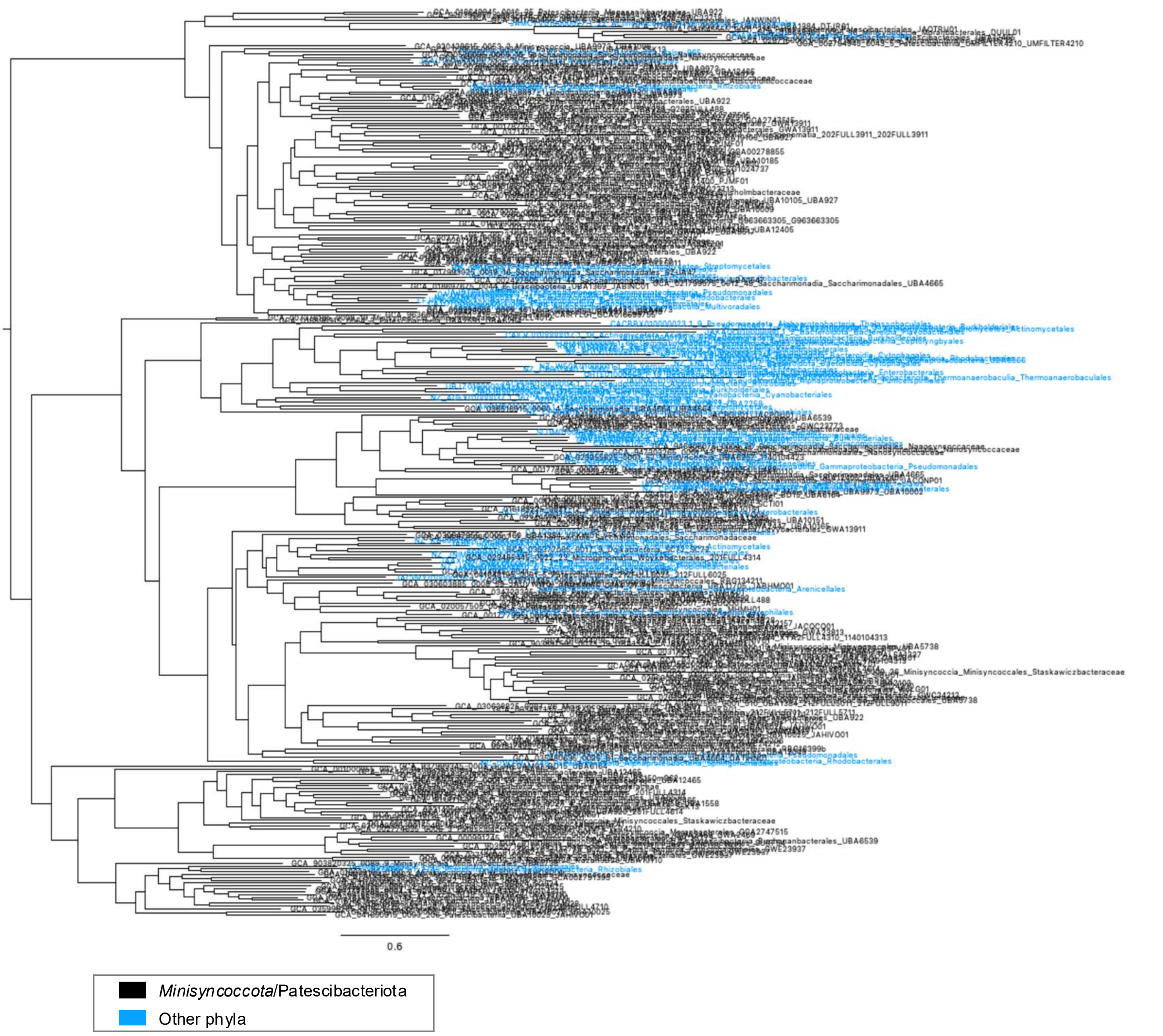


**Supplementary Fig. S15 | Phylogenetic trees of enzymes involved in nucleotide synthesis showing horizontal transfer of these genes.**

**f**, Maximum-likelihood tree estimated from an alignment of pyrimidine biosynthesis protein aspartate dihydroorotase (COG0044). (continued on next page)

**g**

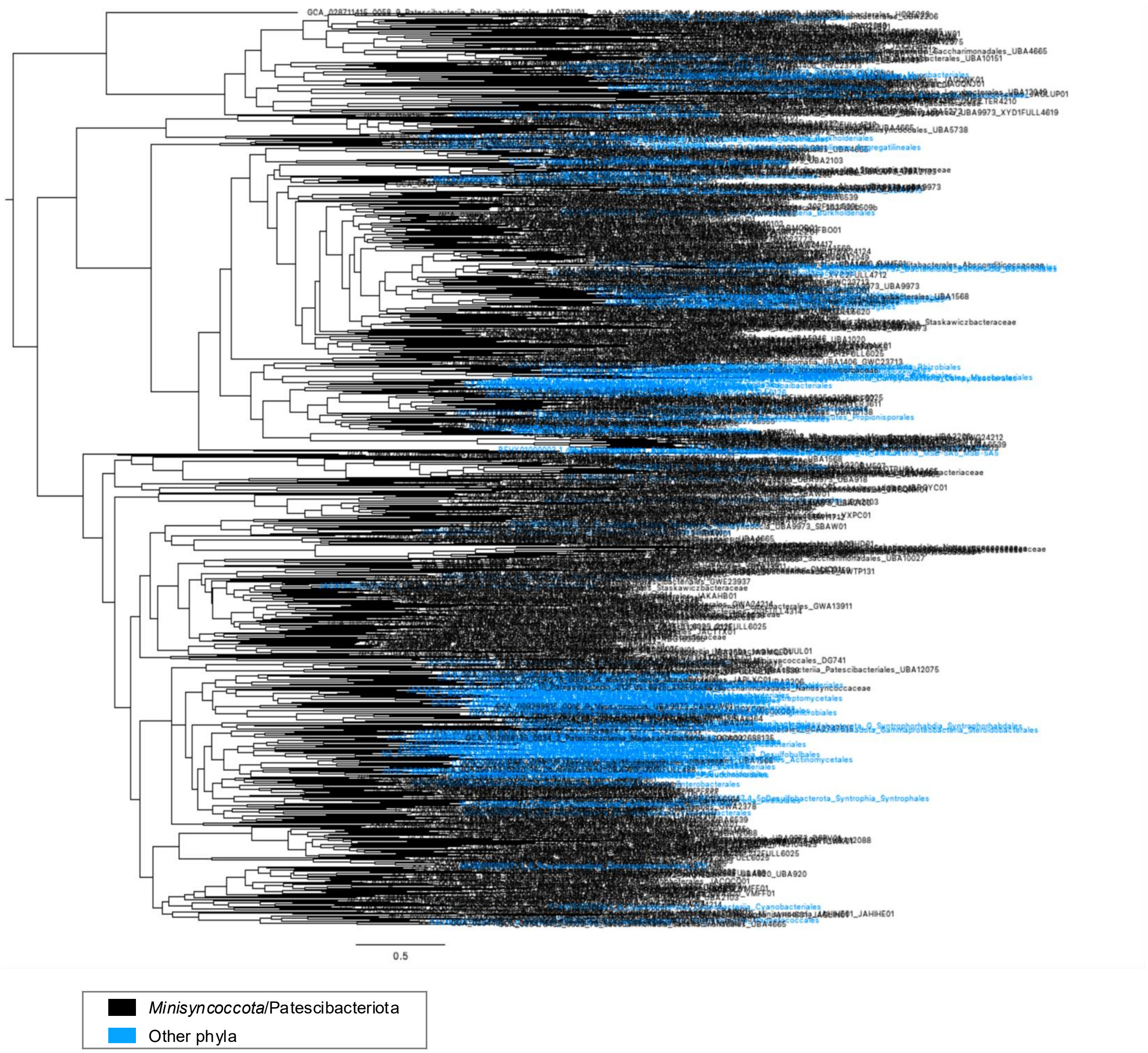


**Supplementary Fig. S15 | Phylogenetic trees of enzymes involved in nucleotide synthesis showing horizontal transfer of these genes.**

**g**, Maximum-likelihood tree estimated from an alignment of pyrimidine biosynthesis protein orotidine-5'-phosphate decarboxylase. (continued on next page)

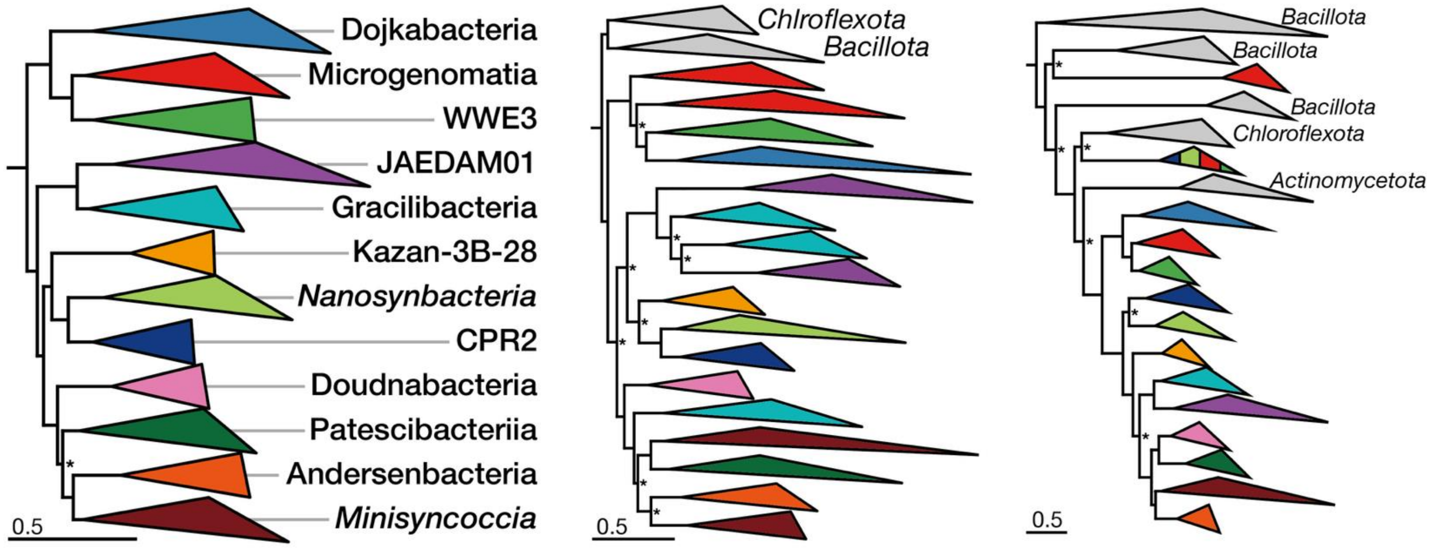


**Supplementary Fig. S16 | Phylogenetic clustering of PNPase and VirB4 in *Minisyncoccota*/Patescibacteriota.**

Maximum-likelihood phylogenies inferred from a concatenated alignment of conserved marker proteins (left), polynucleotide phosphorylase (PNPase; middle), and the type IV secretion system ATPase VirB4 (right). Colors indicate *Minisyncoccota*/Patescibacteriota class-level lineages, and gray indicates sequences from other *Bacillati* phyla. Asterisks indicate nodes with ultrafast bootstrap approximation support <95% or SH-like approximate likelihood ratio test support <80%; all other unlabeled nodes had support values above these thresholds.

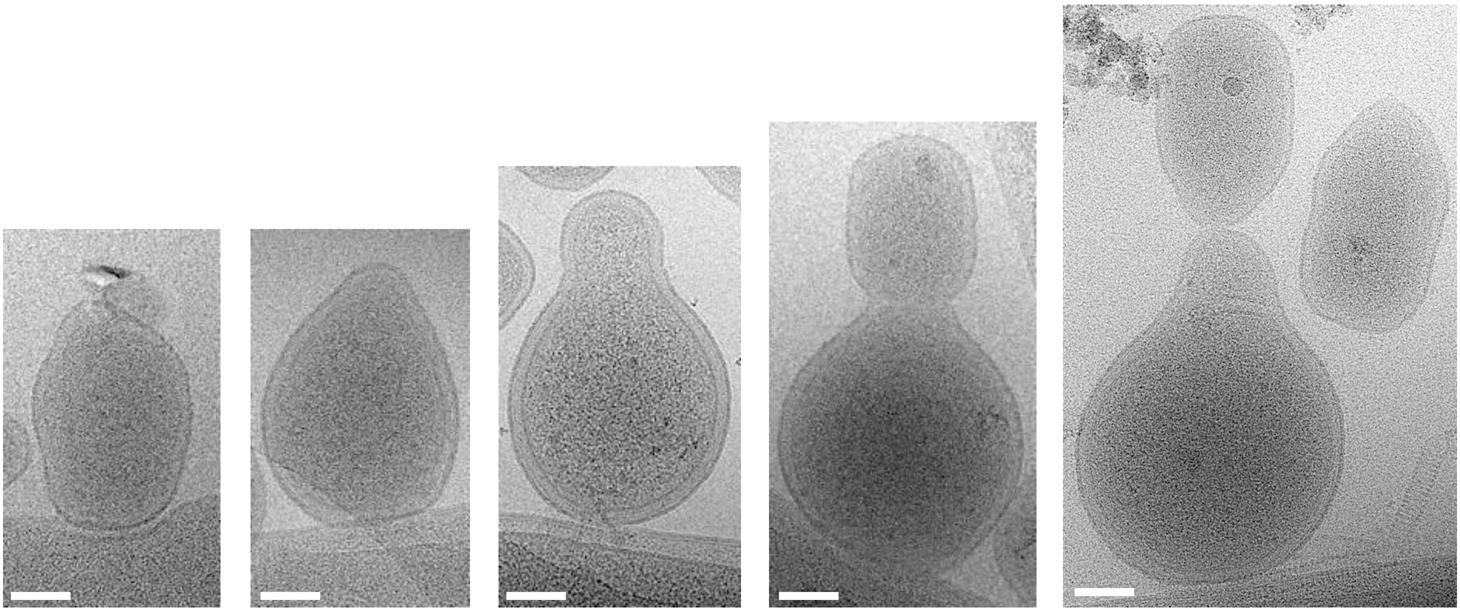

**Supplementary Fig. S17 | Cryo-EM visualization of OT8 enlargement and polar division on the Mc4 surface.**
Representative cryo-electron microscopy images of OT8 cells attached to Mc4, showing cell enlargement and morphological remodeling associated with polar division of OT8. Scale bars, 100 nm.

**a**

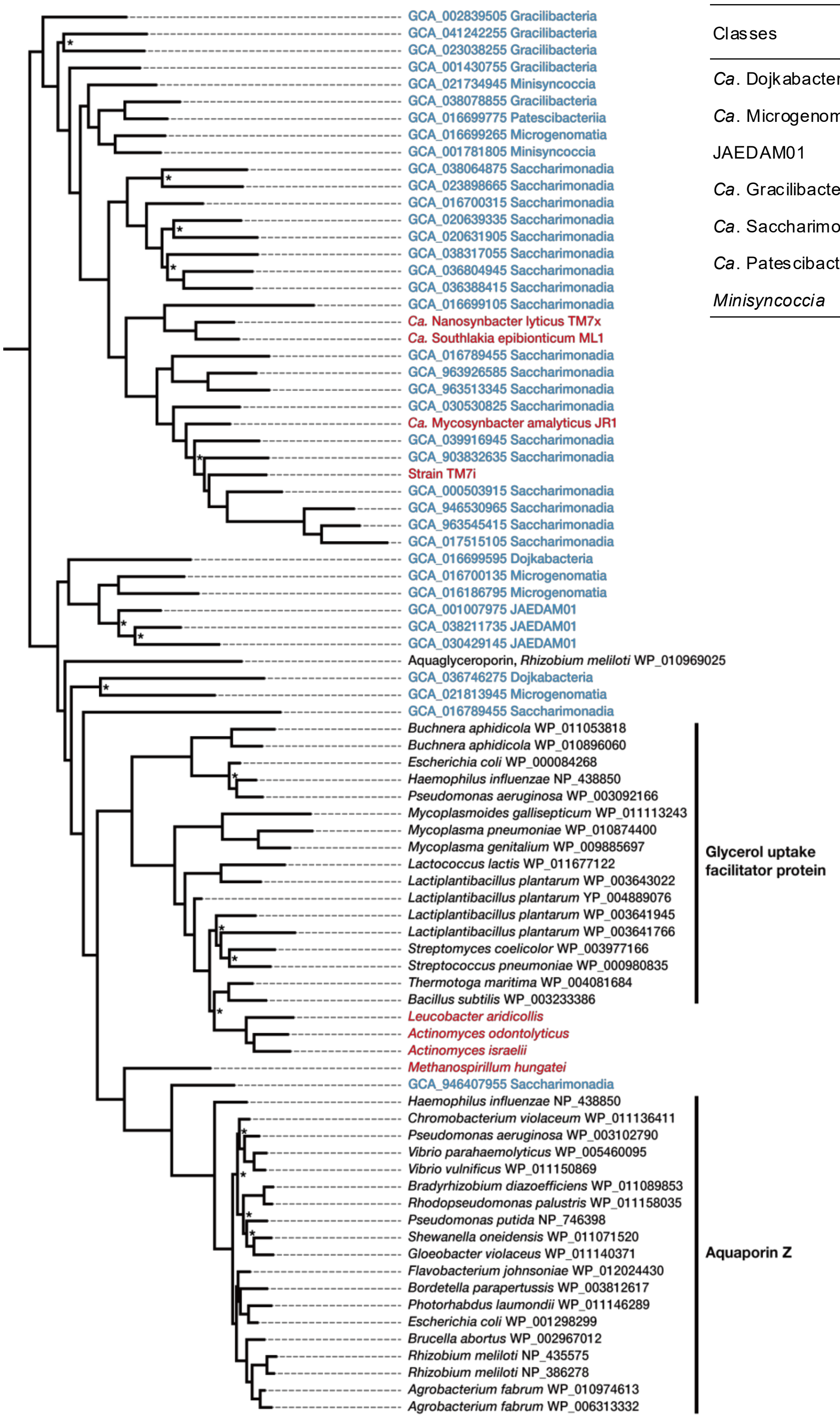


**b**

| Classes | # of MAGs | MIP | Aqp-like |
|---|---|---|---|
| *Ca*. Dojkabacteria | 100 | 4 | 0 |
| *Ca*. Microgenomatia | 624 | 48 | 0 |
| JAEDAM01 | 169 | 26 | 11 |
| *Ca*. Gracilibacteria | 256 | 78 | 1 |
| *Ca*. Saccharimonadia | 1050 | 415 | 48 |
| *Ca*. Patescibacteriia | 737 | 9 | 0 |
| *Minisyncoccia* | 2195 | 57 | 6 |

**Supplementary Fig. S18 | Phylogeny and distribution of major intrinsic proteins (MIPs) in *Minisyncoccota*/Patescibacteriota.**
**a**, Maximum-likelihood tree of protein sequences annotated as major intrinsic proteins (MIPs). Sequences identified by Conserved Domain Search are shown for previously cultured CPR members and their hosts (red), other *Minisyncoccota* lineages (blue), and UniProtKB reviewed MIPs (black). Asterisks denote nodes with ultrafast bootstrap approximation and SH-like approximate likelihood ratio test replication values < 95% and < 80%, respectively. Protein sequences of the formate channel FocA, the nitrite transporter NirC, and the inner membrane protein YfdC from *Escherichia coli* were used as outgroups. **b**, Number of metagenome-assembled genomes (MAGs) from each CPR class that possess annotated MIPs and aquaporin (Aqp)-like protein.

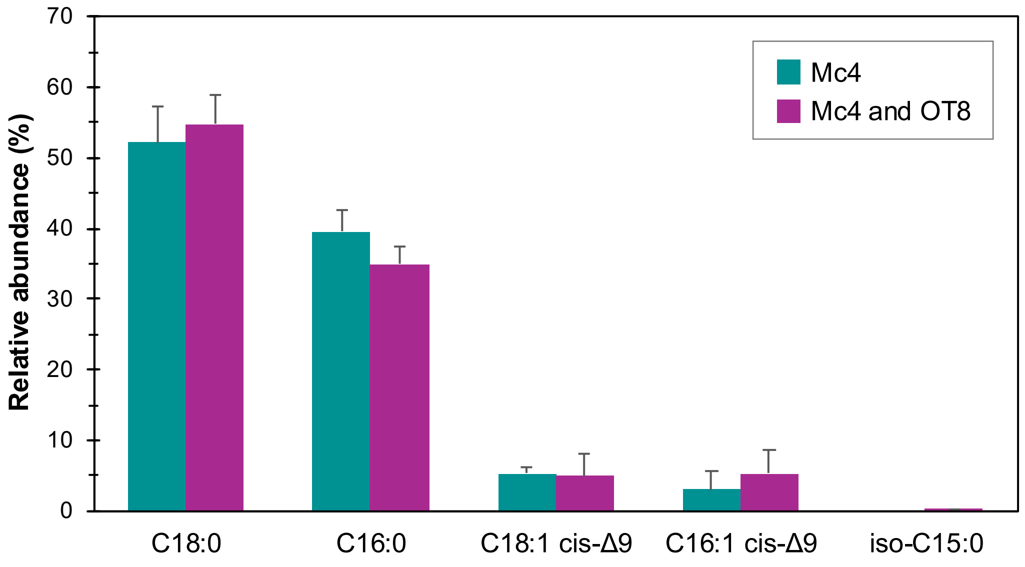


| | Mc4 | | | Mc4 and OT8 | | |
|---|---|---|---|---|---|---|
| | 1 | 2 | 3 | 1 | 2 | 3 |
| C18:0 | 46.08 | 55.07 | 55.21 | 57.97 | 50.4 | 56.13 |
| C16:0 | 42.32 | 39.87 | 36.46 | 36.89 | 32.07 | 36.02 |
| C18:1 cis-Δ9 | 6.41 | 5.06 | 4.58 | 2.34 | 8.43 | 3.77 |
| C16:1 cis-Δ9 | 5.19 | 0 | 3.74 | 2.81 | 9.1 | 4.08 |
| iso-C15:0 | 0 | 0 | 0 | 0.33 | 0 | 0.18 |

**Supplementary Fig. 19 | Cellular fatty acid composition of axenic Mc4 cultures and Mc4–OT8 co-cultures.**

Relative abundances of fatty acids detected in cells harvested from anoxic axenic Mc4 cultures and Mc4–OT8 co-cultures during the late exponential growth phase are shown. Bar plots represent mean values, and error bars indicate standard deviation calculated from three independent biological replicates, each derived from separately grown cultures. Individual replicate values are provided in the table below. A minor branched-chain fatty acid, iso-C15:0, was detected exclusively in Mc4–OT8 co-cultures at very low abundance. Notably, the Mc4 genome encodes FabH (β-ketoacyl-ACP synthase III), a key enzyme required for branched-chain fatty acid biosynthesis, whereas no homolog of FabH is present in the OT8 genome.
